# The maize preligule band is subdivided into distinct domains with contrasting cellular properties prior to ligule outgrowth

**DOI:** 10.1101/2023.02.06.527228

**Authors:** Wesley Neher, Carolyn G. Rasmussen, Siobhan A. Braybrook, Vladimir Lažetić, Claire E. Stowers, Paul T. Mooney, Anne W. Sylvester, Patricia S. Springer

## Abstract

The maize ligule is a fringe of epidermis-derived tissue, which arises from the preligule band (PLB) at a boundary between the blade and sheath. A hinge-like auricle also develops immediately distal to the ligule and contributes to blade angle. Here, we characterize the stages of PLB and early ligule development in terms of topography, cell area, division orientation, cell wall rigidity, and auxin response dynamics. Differential thickening of epidermal cells and localized periclinal divisions contributed to the formation of a ridge within the PLB, which ultimately produces the ligule fringe. Patterns in cell wall rigidity were consistent with the subdivision of the PLB into two regions along a distinct line positioned at the nascent ridge. The proximal region produces the ligule, while the distal region contributes to one epidermal face of the auricles. Whereas the auxin transporter PIN1 accumulated in the PLB, observed differential auxin transcriptional response did not underlie the partitioning of the PLB. Our data demonstrate that two zones with contrasting cellular properties, the preligule and preauricle, are specified within the ligular region prior to ligule outgrowth.

**Summary Statement:** Changes in cell geometry, division orientation, and cell wall mechanics underlie maize ligule morphogenesis. The establishment of mechanically distinct epidermal domains coincides with topographical changes during early ligule outgrowth.

## Introduction

Organogenesis in plants is dependent on positionally determined cell fate patterning together with regulation of cell division and expansion. Morphogenesis and differentiation give rise to diverse leaf shapes with distinct domains, such as petiolate leaves in many eudicots and sheathing leaves in grasses (Moon and Hake, 2011). Early morphogenesis of these leaf domains involves changes in cell growth. The rate and orientation of cell divisions relative to cell expansion therefore play an important role in sculpting leaf shape (Echevin et al., 2019). How cell division patterns and cell expansion are regulated remains an open question and is critical to understand how leaf domains develop.

The mechanisms controlling leaf initiation are relatively well-understood. In the shoot apical meristem (SAM), polar auxin transport by PIN-FORMED (PIN) proteins directs auxin accumulation, which helps specify leaf anlagen (Benková et al., 2003; Okada et al., 1991). Early events associated with leaf initiation include a local maximum of auxin transcriptional responses (Gallavotti et al., 2008; Heisler et al., 2005), downregulation of *Class I KNOTTED1-LIKE HOMEOBOX* (*KNOXI*) genes (Hake et al., 2004; Lin et al., 2003; Ori et al., 2000), periclinal divisions in the L2 (Medford et al., 1992; Vaughan, 1955), and cell wall loosening in subepidermal cell layers (Peaucelle et al., 2011), which all precede leaf primordium emergence. A distinct boundary domain is established between the meristem and an incipient leaf primordium, where cell division and expansion are repressed, which helps facilitate the separation of the organ from the meristem (Hussey, 1971; Kwiatkowska and Dumais, 2003). During the emergence of leaf primordia, organ polarity can be established along three perpendicular axes: transverse, from adaxial (facing the meristem) to abaxial (facing away from meristem), mediolateral, and proximodistal. Cell fate, as well as differences in the rate and direction of cell division and expansion, are patterned along these axes. The boundary is maintained at the base of the organ throughout development and can be recapitulated at other locations on the developing organ (Bouré et al., 2022; Johnston et al., 2014; Nahar et al., 2012; Xiao et al., 2022).

Boundary domains are essential to specify proper morphogenesis and contain distinct cells with altered signaling and cell wall properties. Mutations in genes encoding boundary-defining transcription factors such as *CUP-SHAPED COTYLEDON2* (*CUC2*), *CUP-SHAPED COTYLEDON3* (*CUC3*), *LATERAL ORGAN BOUNDARIES* (*LOB*), and *LATERAL ORGAN FUSION1* (*LOF1*) lead to improper organ separation due in part to derepression of cell division and expansion within the boundary domain (Bell et al., 2012; Gendron et al., 2012; Hibara et al., 2006; Lee et al., 2009). While mutant studies highlight the importance of boundary-defining transcription factors, the mechanisms regulating cell growth in boundaries are not understood. LOB regulates brassinosteroid (BR) catabolism in boundary domains as one mechanism of limiting growth (Arnaud and Laufs, 2013). Cell wall-modifying genes are enriched among the transcriptional targets of BRs, and BR signaling is known to affect cell wall composition and structure, suggesting that cell wall biophysical properties are a component of boundary function (Bai et al., 2012; Graeff et al., 2021; Sun et al., 2010). Consistent with this, cell wall-related gene ontology terms are also significantly enriched among the transcriptional targets of LOB, while CUC2 represses many genes associated with cell wall loosening (Bell et al., 2012; Bouré et al., 2022; Cucinotta et al., 2018). Other experiments show that cell wall-related genes are enriched among highly translated transcripts in the boundary (Tian et al., 2014). Additionally, cells in boundary domains have more rigid cell walls, as measured with atomic force microscopy (AFM) (Bouré et al., 2022; Sampathkumar et al., 2019). Changes in cell wall composition or remodeling activity could contribute to the decreased rate of cell expansion in boundary domains, but more experiments are needed to determine how the boundary function modulates growth. While the SAM-leaf primordium boundary has been relatively well-studied, less is known about how other developmental boundaries are specified and contribute to organogenesis.

A challenge in analyzing the leaf-SAM boundary is its physical inaccessibility. The maize leaf provides a unique opportunity to study an accessible boundary at the ligular region, which plays an important role in the proximodistal patterning of the leaf. The two largest domains of grass leaves are the proximal sheath and the distal blade separated by the ligular region (Fig. S1), where several specialized structures develop. At the blade-sheath junction, a thin epidermis-derived flap called the ligule covers the gap between consecutive ensheathing leaves. Also at the blade-sheath junction, two wedge-shaped structures called auricles develop on both sides of the midrib. Auricles are thought to facilitate the outward bending of the blade to optimize photosynthetic light capture (Emerson, 1912). The ligule is derived from a distinct region of the adaxial epidermis called the preligule band (PLB), a narrow, linear boundary domain between the preblade and presheath of the leaf primordium (Becraft et al., 1990; Sylvester et al., 1990). Due to physical proximity and the genetic links between the PLB, ligule and auricle, the adaxial epidermal portion of the pre-auricle is also hypothesized to be derived from the PLB and/or from blade tissue adjacent to the upper boundary of the PLB. These hypotheses have not yet been resolved.

Factors involved in the development of the ligular region have been genetically identified, including the maize transcription factors *liguleless1* (*lg1*) and *liguleless2* (*lg2*), which define the blade-sheath boundary and specify ligule and auricle development in a partially redundant manner (Becraft et al., 1990; Walsh et al., 1998). Single mutants *lg1-R* and *lg2-R* affect the position of the blade-sheath boundary and alter the pattern of ligule and auricle development, while the *lg1-R*; *lg2-R* double mutant has an indistinct blade-sheath boundary and lacks both ligule and auricle (Foster et al., 2004; Harper and Freeling, 1996). Mutations in *liguleless* genes result in more vertical leaf angles because the auricles do not develop properly (Emerson, 1912). Rice *lg1* and *lg2* mutants display mutant phenotypes similar to those in maize, suggesting functional conservation in grasses, despite differences in ligular region structures (Lee et al., 2007; Wang et al., 2021).

Changes in cell division and expansion are observed in development of the ligule (Becraft et al., 1990; Sharman, 1942; Sylvester et al., 1990). Cells divide more frequently in the adaxial epidermis in the PLB based on the emergence of new cross-walls (Freeling, 1992; Sylvester et al., 1990). *lg1*transcript accumulates at this site of increased division (Johnston et al., 2014; Moreno et al., 1997). Several rounds of epidermal anticlinal divisions, along with decreased cell expansion, reduce cell surface area (Becraft et al., 1990; Sylvester et al., 1990). The PLB becomes visible as a narrow band of small cells running laterally across the adaxial epidermis between the blade and sheath domains. After several rounds of anticlinal divisions, periclinal divisions are observed in both the PLB and the underlying ground tissue, and a ridge forms within the PLB (Sylvester et al., 1990). The auricle differentiates between the ridge and the blade while the ligule develops as a fringe of cells growing up and out from the PLB ridge (Freeling, 1992; Sylvester et al., 1990).

The proximodistal transcriptomic profile of the ligular region has been analyzed with high spatial resolution by laser-capture microdissection followed by RNA-seq in both wild-type B73 and *lg1-R* mutants (Johnston et al., 2014). This revealed that several genes involved in leaf initiation and patterning at the SAM are redeployed later during ligule development. Notably, transcript levels of *KNOXI* class and other boundary-associated transcription factor genes such as *CUP-SHAPED COTYLEDON2-like* (*CUC2-like*) are significantly higher in the PLB (Johnston et al., 2014). *In situ* hybridization showed *CUC2-like* transcripts were detected throughout the PLB early in development, but later became restricted to the distal zone of the PLB, where a cleft will form as the ligule grows out. Xiao et al. (2022) further supported the link between *liguleless2* and boundary-associated gene expression in the context of bract suppression in the inflorescence. These patterns of gene expression support the idea that the PLB functions as a boundary domain. While the SAM/leaf boundary is characterized by a low mitotic rate (Hussey, 1971), the PLB displays increased cell division relative to neighboring regions (Becraft et al., 1990; Sylvester et al., 1990). However, reduced cell size is a common feature between the PLB and SAM/leaf boundary (Becraft et al., 1990; Hussey, 1971). Further, PIN-like proteins and several auxin-responsive genes are upregulated in the PLB, suggesting a role for auxin in ligule development (Johnston et al., 2014; Moon et al., 2013). One proposed model is that *PIN-like*, *KNOXI*, and *CUC2-like* genes are expressed in the early PLB, but subsequent antagonism by auxin responses restricts the expression of boundary-associated genes to the cleft, resulting in further refinement of the PLB into subdomains (Johnston et al., 2014).

Here we document that the stages of ligule development correlate with sheath length, providing a convenient proxy for estimating ligule developmental stage. During the early stages of ligule outgrowth, we compared cell depth, division orientation, and cell wall rigidity along the proximodistal axis. There was a clear divergence in cellular properties between proximal and distal PLB-derived domains prior to ligule outgrowth. Hypothesizing that auxin dynamics may underlie this differentiation, we examined the accumulation of auxin reporters during ligule development. While the auxin transporter PIN1a was observed in the PLB, we did not detect local differences in auxin transcriptional responses prior to ligule outgrowth. Our findings of cell growth patterns and biophysically distinct regions within the PLB may be explained by structural remodeling of cells required for the establishment and physical separation of a new growth axis.

## Results

### Ligule developmental stages correlate with sheath length

To establish developmental reference stages for ligule morphogenesis, we characterized features such as topography and cell size. Existing literature describes the stages of ligule development relative to plastochron number, a value indicating the relative age of a leaf. Plastochron number is difficult to determine because it requires either sectioning or complete dissection down to the meristem (Johnston et al., 2014; Sylvester et al., 1990). In contrast, we used sheath length as a reliable and convenient proxy for predicting the stage of ligule development. Stages of ligule development were characterized in relation to sheath length in two-, three-, and four-week-old plants (Figs. 1, S2). Sheath lengths were measured in sequentially dissected leaves and ligule regions were observed by scanning electron microscopy (SEM; Fig. 1A-E). Ligule developmental stages were also visualized using confocal microscopy of leaves expressing YFP-TUBULIN (Fig. 1F-J). Although the ligule develops continuously and progressively, distinct morphological stages of ligule development correlated significantly with sheath length in expanding adult leaves (Fig. 1K).

**Figure 1:**
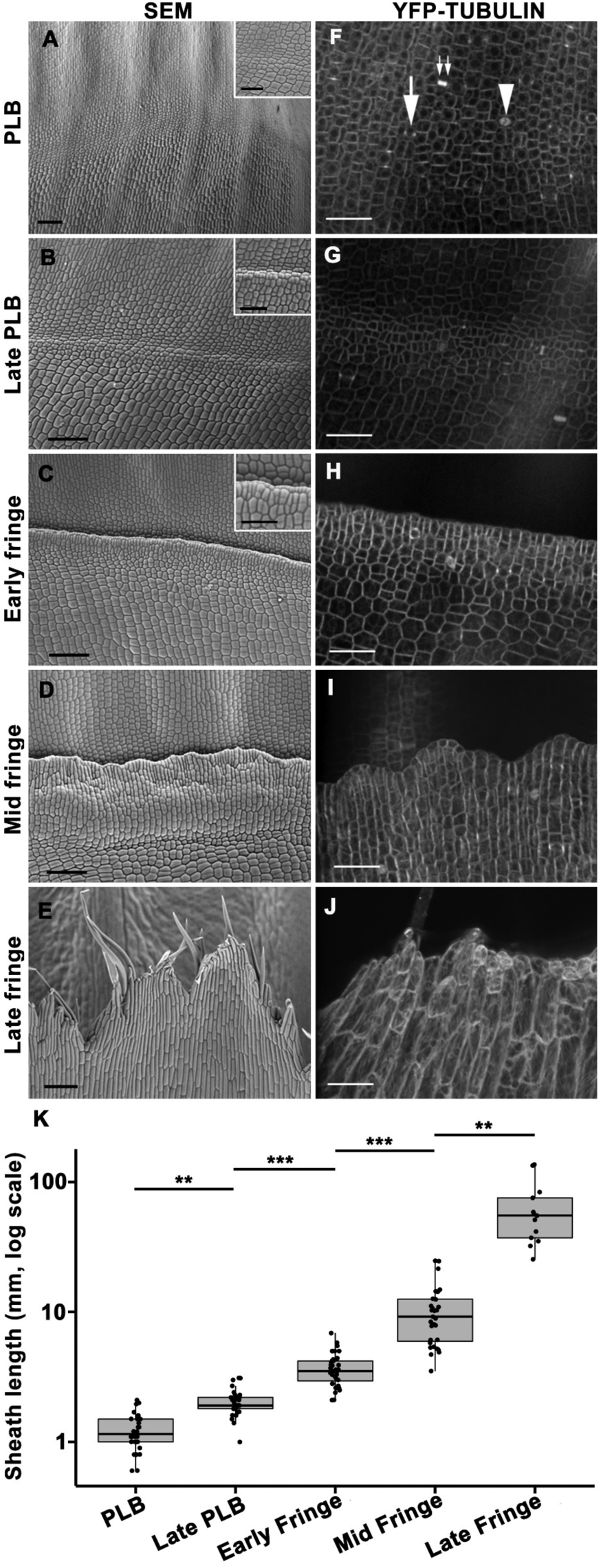
Stages of ligule development correlate with sheath length. (A-E) Scanning electron micrographs show stages of ligule development in four-week old plants. In these images, the sheath is positioned towards the bottom, with the blade at the top. Scale bar = 100 μm. Higher magnification insets in (A-C) show small cells in the preligule band, scale bar = 50 μm. (F-J) YFP-TUBULIN marker shows the division figures used for analysis including (F) preprophase bands (arrow), mitotic spindles (arrowhead) and phragmoplasts (double arrow). Mid and late fringe micrographs are maximum projections of Z stacks three to five µm in depth. Scale bar = 50 μm (K) Box-whisker plot showing sheath height correlates with ligule stage. The box indicates the 25% and 75% quartile, the median is indicated by a bar, while whiskers indicate the range of data. N = 32 (PLB), 30 (Late PLB), 35 (Early Fringe), 33 (Mid Fringe), 13 (Late Fringe). Statistical significance assessed via Kruskall-Wallis test with Dunn’s post-hoc and Benjamini-Hochberg p-value adjustment for multiple comparisons. Only comparisons between consecutive stages are shown here. ** indicates p<0.05, *** indicates p<0.01.

The stages distinguishable by SEM were defined as preligule band (PLB), late PLB, and early, mid, and late fringe. At a median sheath length of 1.2 mm, the PLB consisted of a band of small cells spanning ∼60-100 μm in proximal/distal length. This measure is consistent with previously published images of longitudinal sections that show the position of the PLB relative to the SAM (Johnston et al. 2014). PLB cells tended to be more isodiametric than the more elongated blade and sheath cells. At this stage, a slight ridge was often visible at the blade sheath junction, with an inflection point at the PLB (Fig. 1A,F). At the late PLB stage, this ridge was very pronounced with the adjacent sheath surface elevated above the blade surface (Fig. 1B,G). Median sheath length was 1.9 mm, and cell area reached a minimum at this stage (Table S1). Leaves in the early fringe stage had a median sheath length of 3.5 mm. Ligule cells had relatively uniform size and shape and were aligned at the leading edge of the ridge, beginning to grow over the more distal PLB-derived cells (Fig. 1C,H). Cell area increased throughout the development of the fringe (Table S1). Leaves in the mid fringe stage had a median sheath length of 8.7 mm. The ligule appeared “corrugated” and uneven relative to the plane of the leaf (Fig. 1D,I). At a median sheath length of 54.1 mm, the ligule was in the late fringe stage defined by elongate hair-like cells at the leading edge of the ligule, which projected over the developing auricle and blade (Fig. 1E,J). These observations demonstrate that ligule development correlates with sheath growth (Figs. 1K, S2). This shows that sheath length can be used to approximate the developmental stage of the ligule.

### Changes in cell division orientation and expansion are associated with PLB and fringe growth

Alterations in cell expansion and division are important characteristics of the preligule band. We used a live cell marker for microtubules, YFP-TUBULIN (Mohanty et al., 2009), to assess cell area and division plane orientation at each of the stages described above. We calculated the relative frequencies of different divisions by classifying the orientation of preprophase bands and phragmoplasts (Fig. 2A-E). Visualizing microtubule structures enabled us to discern an earlier developmental stage than was visible with SEM, which we termed the early PLB stage. During this stage, the predominance of longitudinal anticlinal divisions (>50%) at the blade-sheath junction was the distinguishing feature, and average cell area was ∼159 µm^2^ (Fig. 2F, Table S1). In the PLB stage, transverse anticlinal divisions were the most frequent and a low frequency of periclinal divisions was observed (Fig. 2F). In addition, the average PLB cell area decreased to ∼135 μm^2^ (Table S1). In the late PLB stage, periclinal divisions were observed most frequently (∼46%) and the average cell area was further reduced to ∼106 μm^2^. These results show reduced cell sizes in the PLB and shifts in division orientation from anticlinal to periclinal by the late PLB stage.

**Figure 2:**
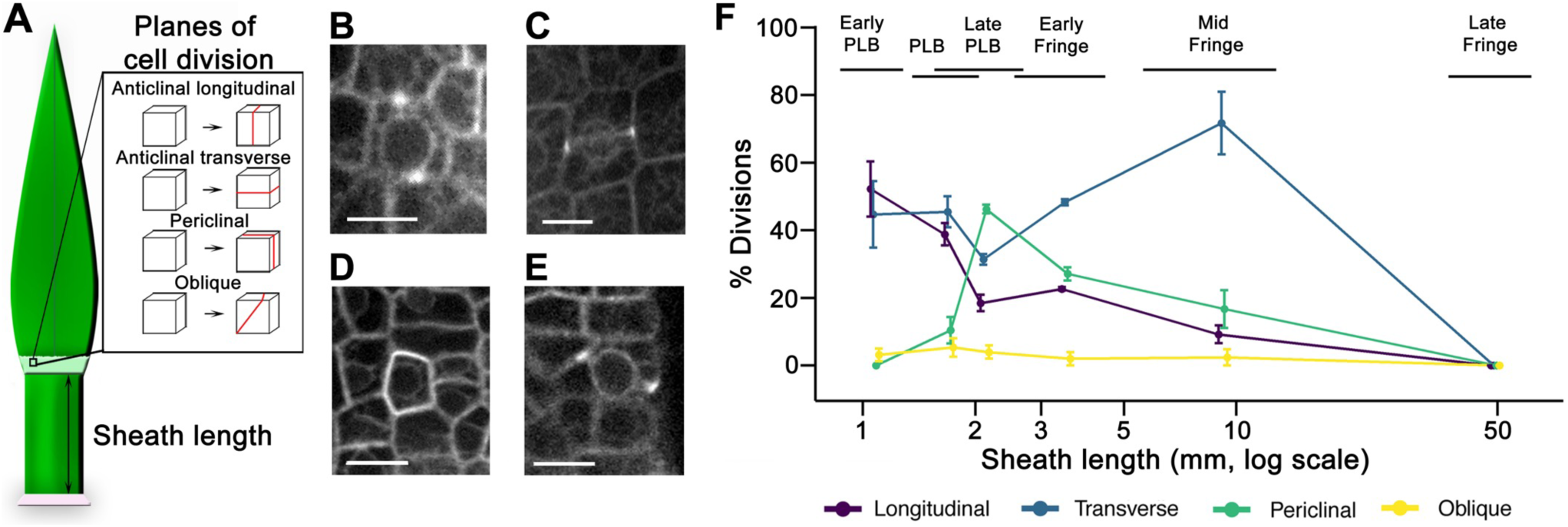
Dynamic changes in division plane orientation during ligule development. (A) Diagram illustrating the position of the PPB, where mitotic cells were imaged. Division plane orientation is illustrated with the red line indicating the new plane of division. (B-E) Examples of preprophase bands in cells expressing YFP-TUBULIN (B, longitudinal; C, transverse; D, periclinal; E, oblique) Scale bars = 15 μm. (F) The percentage of dividing cells that exhibit each division orientation at each stage of ligule development. Error bars indicate standard error. N=3-5 leaves per stage, 11-46 mitotic cells per leaf. Black horizontal bars indicate the range of sheath lengths for the leaves that were measured at each stage. Color lines are included to help visualize the trend for each division type, and do not imply any relationship between samples at each stage.

During the early fringe stage, periclinal divisions were reduced, with ∼48% of the divisions oriented in the transverse anticlinal plane. Cell expansion increased so that early fringe cells were ∼80% larger than late PLB cells. Mid fringe cells divided mostly in the transverse anticlinal orientation (71%), and cell area increased dramatically. Ligule cells in the late fringe stage no longer divided, but continued expanding, producing larger and more variably sized cells (Fig. 2F and Table S1). These results show that spatially constrained divisions initially produce the ligule fringe, with cell expansion driving growth in the later stages.

### Differential cell expansion and division orientation within the PLB contribute to the formation of the preligule ridge

During early ligule development, a ridge forms so that the sheath surface is elevated relative to the blade surface. While periclinal divisions in the PLB and underlying mesophyll are known to contribute to the formation of this ridge, proximodistal differences in cell thickness (depth) have not been quantified. To determine whether differential thickening of epidermal cells contributed to the formation of the ridge, we measured cell depth in the epidermis of the sheath, ligular region, and blade during the early, mid, and late stages of PLB development (Fig. 3A). In the early PLB stage, cell depth was uniform, averaging 11-12 μm in the sheath, PLB, and blade (Fig. 3A). During the PLB stage, the sheath and proximal ligular region cells averaged around 16.4 μm deep, while the distal ligular region and blade cells were around 13 μm deep (Fig. 3A). This relative thickening of the sheath coincided with the formation of the ridge. By the late PLB stage, the rate of periclinal divisions in the PLB increased and the preligule ridge became more pronounced, with cells on the proximal side of the ridge averaging 19.2 μm deep (Fig. 3A). Meanwhile, the distal PLB-derived cells were the thinnest in the epidermis, averaging 13.9 μm deep. Confocal projections suggested that differential thickening likely also occurs in underlying cell layers; however, reduced signal prevented measuring cell thickness in deeper layers. Our findings regarding epidermal cell depth are consistent with previously published TEMs of the developing ligule (Sharman, 1941; Sharman, 1942). These data show that differential cell thickening contributes to the changes in epidermal topography during the early stages of ligule development.

**Figure 3:**
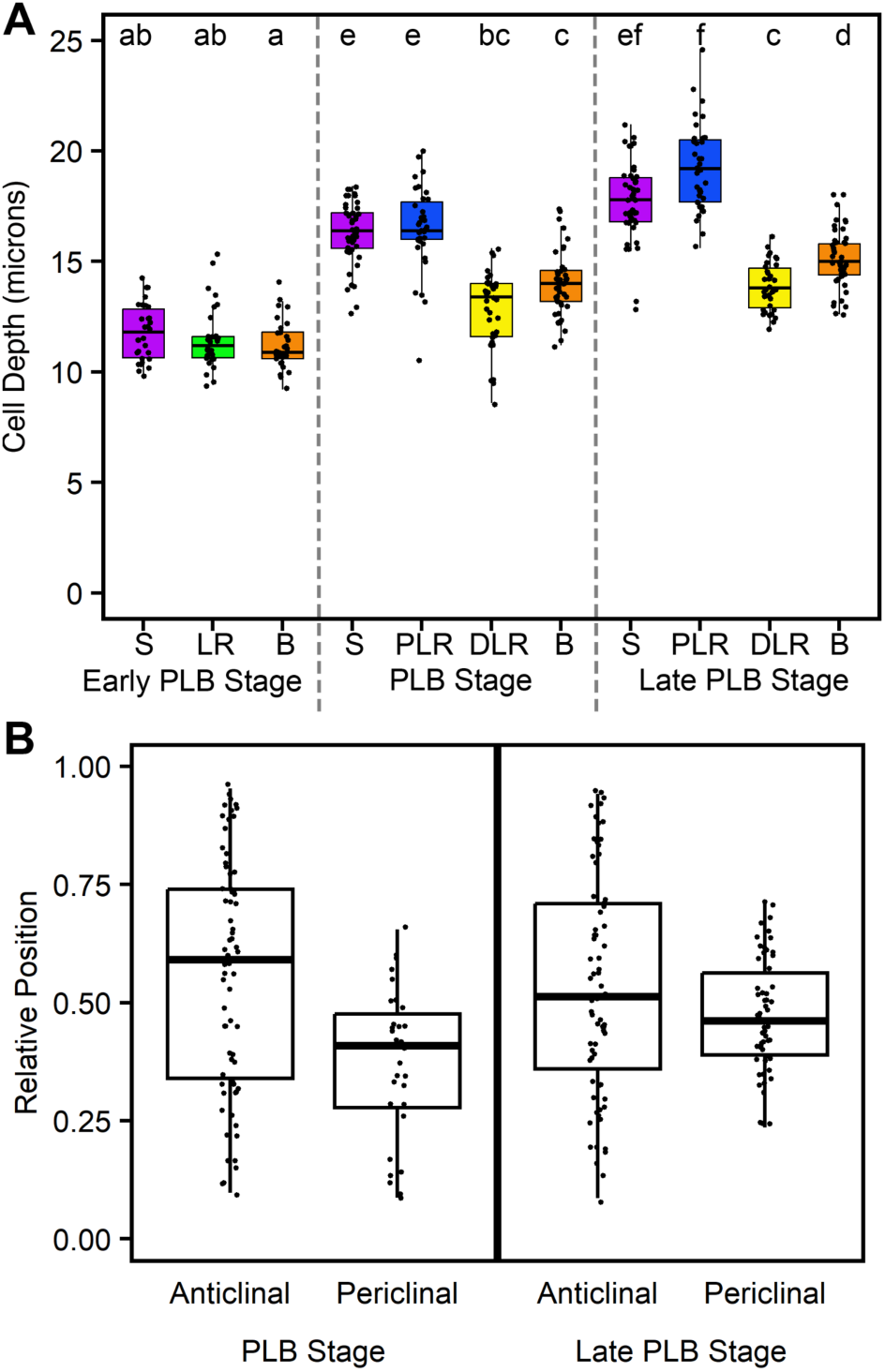
Differential cell thickening and periclinal divisions contribute to formation of the preligule ridge. Cell depth was measured from confocal z-stacks at anticlinal faces of cells that had not yet undergone periclinal divisions, in the ligular region of the maize leaf adaxial epidermis. (n=3 leaves per stage, 10-15 cells per region per leaf) Significance was determined via Kruskal-Wallis test with Dunn’s post-hoc, adjusting p-values via the Benjamini-Hochberg method for multiple comparisons, at an alpha of 0.05. Letter rankings indicate comparisons between all stages and epidermal zones. S=Sheath, LR=Ligular Region, PLR=Proximal Ligular Region, DLR=Distal Ligular Region, B=Blade (A) Relative position of periclinal divisions within the ligular region was determined from confocal micrographs of plants expressing either CFP-Tubulin or TAN-YFP. Position 0 is the proximal extremity of the ligular region while 1 is the distal extremity.

While periclinal divisions are known to drive early ligule outgrowth from the PLB, it is not clear whether periclinal divisions occur throughout the whole ligular region or are specific to a subset of cells that form the ligule. The relative positions of periclinal and anticlinal divisions within the ligular region were determined using either YFP-TUBULIN or TANGLED-YFP, a protein which localizes to the division site (Martinez et al., 2017; Walker et al., 2007). The ligular region was defined as the zone of reduced cell area between the blade and sheath (Fig. S3). In the PLB stage, sporadic periclinal divisions were visualized in the proximal 2/3^rds^ of the ligular region but not in the distal 1/3^rd^ (Fig. 3B). In the late PLB stage, periclinal divisions were exclusively observed in the median 50% of the ligular region, localized to the nascent preligule ridge, but absent from both extremities (Fig. 3B). At both stages, anticlinal divisions were broadly distributed over the entire proximodistal length of the ligular region (Fig. 3B). Therefore, epidermal periclinal divisions occur in the proximal ligular region, but not in the distal cells that contribute to the auricles.

### Mechanical changes within the epidermis precede ligule outgrowth

Differences in cell size and division orientation indicated that two zones with contrasting cellular behavior are established in the ligular region prior to emergence of the fringe. The elastic properties of the cell wall often correlate with cell expansion and reflect physical differences both between different cell populations and subcellular cell wall domains (Bou Daher et al., 2018; Peaucelle et al., 2011). We sought to identify cell wall mechanical patterns in epidermal cells during development of the ligular region. Atomic force microscopy (AFM) uses a physical probe to measure the topography and various physical characteristics of surfaces. Raw AFM indentation data can be processed using a Hertzian contact model to calculate the indentation modulus (IM) of a surface. IM is the complex elastic (reversible) stiffness of the area being indented, or the relationship between the pressure applied to a surface and the resulting elastic area change of that surface. High IM values indicate greater rigidity of a surface because more pressure is necessary to deform the surface. Hereafter, IM will be referred to as rigidity. To eliminate turgor pressure and only consider cell wall mechanical properties, leaves were plasmolyzed prior to being measured.

We used AFM to measure the rigidity of cell walls across epidermal regions in B73 leaves from the early PLB stage through the early fringe stage. Periclinal walls had relatively low rigidity, while anticlinal walls had higher rigidity, consistent with previous experiments in plasmolyzed tissue (Bou Daher et al., 2018; Peaucelle et al., 2011; Sampathkumar et al., 2019). To reveal tissue-scale patterns in rigidity along the proximodistal axis, we analyzed the AFM scans using a sliding window approach (see methods, Fig. 4). Generally, the sheath had lower average rigidity than the blade at all stages of development. During early PLB and PLB stages, the central ligular region was the most rigid epidermal zone (Fig. 4A,B). During late PLB and early fringe stages, a different mechanical pattern was observed in the ligular region. A distinct “transition” zone at the proximal end of the ligular region contained cells that were similar in shape to the sheath cells, but smaller and mechanically softer. The center of the ligular region had very small cells with the lowest average rigidity. The distal ligular region, meanwhile, remained the most rigid epidermal zone (Fig. 4C,D). Similar results were obtained in Mo17 leaves (Fig. S4), indicating that these patterns were not unique to the B73 genetic background. Therefore, a transition in mechanical properties occurs between the PLB and late PLB stages, with significant softening in the middle of the PLB, while the distal pre-auricle region remains rigid.

**Figure 4:**
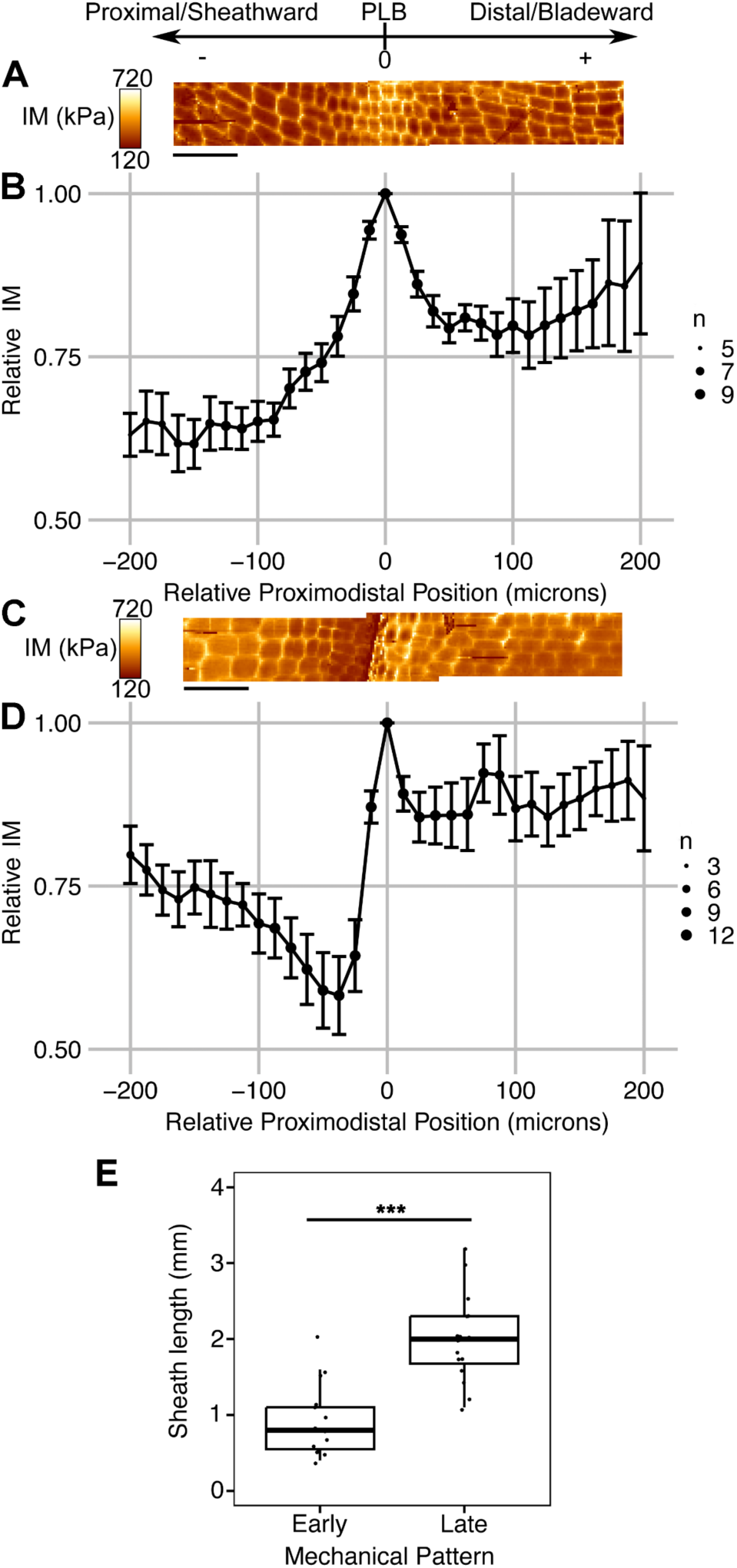
AFM analysis of ligular region reveals two distinct mechanical phases during PLB development. Scale at top shows orientation of leaf for AFM scans and sliding window analysis, with the local maximum in IM for each leaf set as position 0. (A) Representative scans of leaf in the PLB stage reveals a local maximum in IM within the PLB region. Color scale indicates the indentation modulus (IM, kPa) for an indentation, with each pixel representing one indentation. Two overlapping 50 x 200 μm scans are shown. (B) Sliding window analysis averaging all B73 samples exhibiting the early pattern (Early PLB and PLB stages; n=9). (C) Representative scans of leaf in the early fringe stage (D) Sliding window analysis averaging all B73 samples exhibiting the late pattern (Late PLB and Early Fringe stages, n=13). Samples included in (D) had a significantly higher sheath length compared to samples in (B), showing that the apparent softening of PLB cells occurs at a later developmental stage. (A,C) are to scale relative to (B,D) Scale bars = 50 μm. Size of dot (n) indicates coverage at that relative position. Error bars = S.E. (E) Sheath lengths of leaves with the early and late mechanical patterns. *** indicates p<0.01, calculated via Mann-Whitney U-test.

To determine when the mechanical changes in the ligular region occur relative to ligule outgrowth, we directly compared topographical features to the rigidity data from the same AFM scans. In PLB stage leaves, a shallow ridge was visible in the ligular region (Fig. S5A,B). Late PLB stage leaves had a steep PLB ridge, but relatively flat blade and sheath regions (Fig. S5C,D). The ligule grows out from this ridge, extending over the pre-auricle cells toward the blade. At early stages, the most rigid cells were centrally located in the PLB, but at later stages were shifted distally, consistent with the position of the nascent ligule-auricle cleft. A distinct low-rigidity band, located on the crest of the ridge, was observed in the late PLB and early fringe stages (Fig. 4D, Fig. S5C,D). Cell wall softening in the late PLB stage precedes the increases in cell area driven by cell expansion during the early fringe stage.

The sliding window method revealed two distinct mechanical patterns in the epidermis, however this analysis may be biased because anticlinal walls are perceived as more rigid than periclinal walls in plasmolyzed tissue (Peaucelle et al., 2011), and cell size varies between epidermal regions. To avoid potential measurement bias due to cell size differences, we manually resampled rigidity from the AFM scans to compare transverse, longitudinal, and periclinal cell wall segments (Fig. S6, Table S2). Generally, manual resampling confirmed the trends reported above, producing rigidity profiles that were similar to the sliding window analyses (Figs. 4, S6A,B, Table S2). In addition, considering each wall orientation separately allowed us to assess elastic asymmetry, defined as differences in the rigidity of different wall orientations (Bou Daher et al., 2018)(Fig. S6C,D, Table S3). For example, in early stage leaves the average rigidity of transverse walls was higher than that of longitudinal walls in the blade and sheath, but not in the PLB, indicating reduced elastic asymmetry in the PLB cells (Fig. S6C, Table S3). The softer longitudinal walls in the blade and sheath are consistent with the primary direction of organ growth. Lastly, this approach enabled us to compare the average rigidity between the early and late stages (Table S4), revealing that the sheath anticlinal walls rigidified significantly in the late stage while the cells on the preligule ridge softened. Manual resampling supported the tissue-level patterns in rigidity observed with the sliding window approach and enabled further comparisons between the two mechanical stages and cell wall segments with different orientations.

### The pattern of PIN1a-YFP signal changes during PLB stages and is ubiquitous in ligule cells

Auxin has roles in many aspects of leaf development, including specification of founder cells (Reinhardt et al., 2003; Scanlon, 2003). Additionally, application of exogenous auxin was sufficient to induce cell wall biochemical and mechanical changes in the SAM (Braybrook and Peaucelle, 2013). Previous work showed PIN auxin-efflux carrier transcripts *PIN1a*, *PIN1c*, *PIN5*, and *SoPIN1/PIN1d/PIN4* accumulate in the PLB, and PIN1a-YFP signal is strong in the PLB and underlying mesophyll (Conklin et al., 2019; Johnston et al., 2014; Moon et al., 2013), suggesting a role for auxin in ligule specification and/or outgrowth (Johnston et al., 2014). Live cell imaging of PIN1a-YFP was conducted during all stages of ligule development. At the early PLB stage, PIN1a-YFP localized to the plasma membrane of cells over the vasculature in both the epidermis and mesophyll (Fig. 5A,B). In the PLB stage, PIN1a-YFP was observed uniformly in the PLB and underlying mesophyll (Fig. 5C,D). The PIN1a-YFP-expressing zone consistently narrowed from ∼60 μm in the PLB stage to ∼45 μm at the late PLB stage, with signal only in the small PLB cells and underlying mesophyll, and not in the cells at the extremities of the ligular region (Fig. 5E,F). In the early fringe stage, the PIN1a-YFP-accumulating zone expanded only in the sheathward/proximal direction (Figs. 5G, 6C). PIN1a-YFP signal was observed in ligule cells at all stages of fringe development (Fig. 5G-I). The narrowing of the zone containing PIN1a-YFP signal correlates with the increase in the periclinal division rate in the late PLB stage and precedes the outgrowth of the ligule fringe.

**Figure 5:**
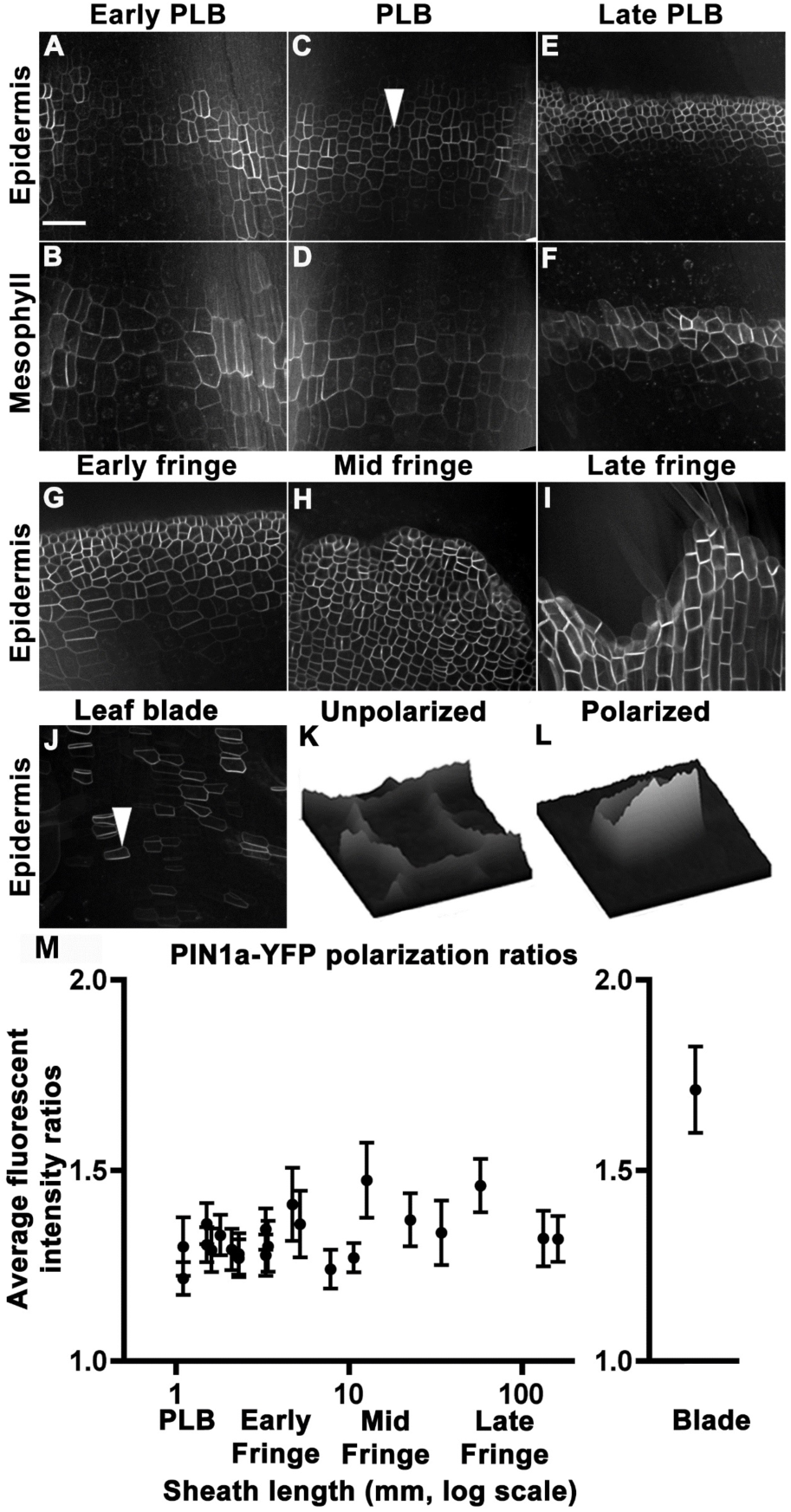
PIN1a-YFP localization in the PLB and ligule. Panels (A,C,E) are single Z plane images of epidermal cells in early PLB, PLB and late PLB respectively; (B,D,F) are corresponding single Z planes of the next cell layer in the subtending mesophyll. PIN1a-YFP was observed in the early fringe (G), mid fringe (H) and late fringe (I). In (J) PIN1-YFP signal was peripheral in blade cells above the ligule fringe. (K,L) Relative fluorescence intensity surface plots are displayed for the cell by arrowheads in (C), showing a PLB cell with low polarity, and in (J), showing a blade cell with increased polarity. (M) PIN1a-YFP polarization ratios compared at ligule stages and in the blade, as calculated from 30 or more cells from four different plants. Each point is the mean polarization ratio for a given sheath height. Error bars are 95% confidence intervals. Blade cells were significantly more polarized than PLB or fringe cells (p value ≤ 0.01 using the Kolmogorow-Smirnov test). Scale bar = 50 μm.

**Figure 6:**
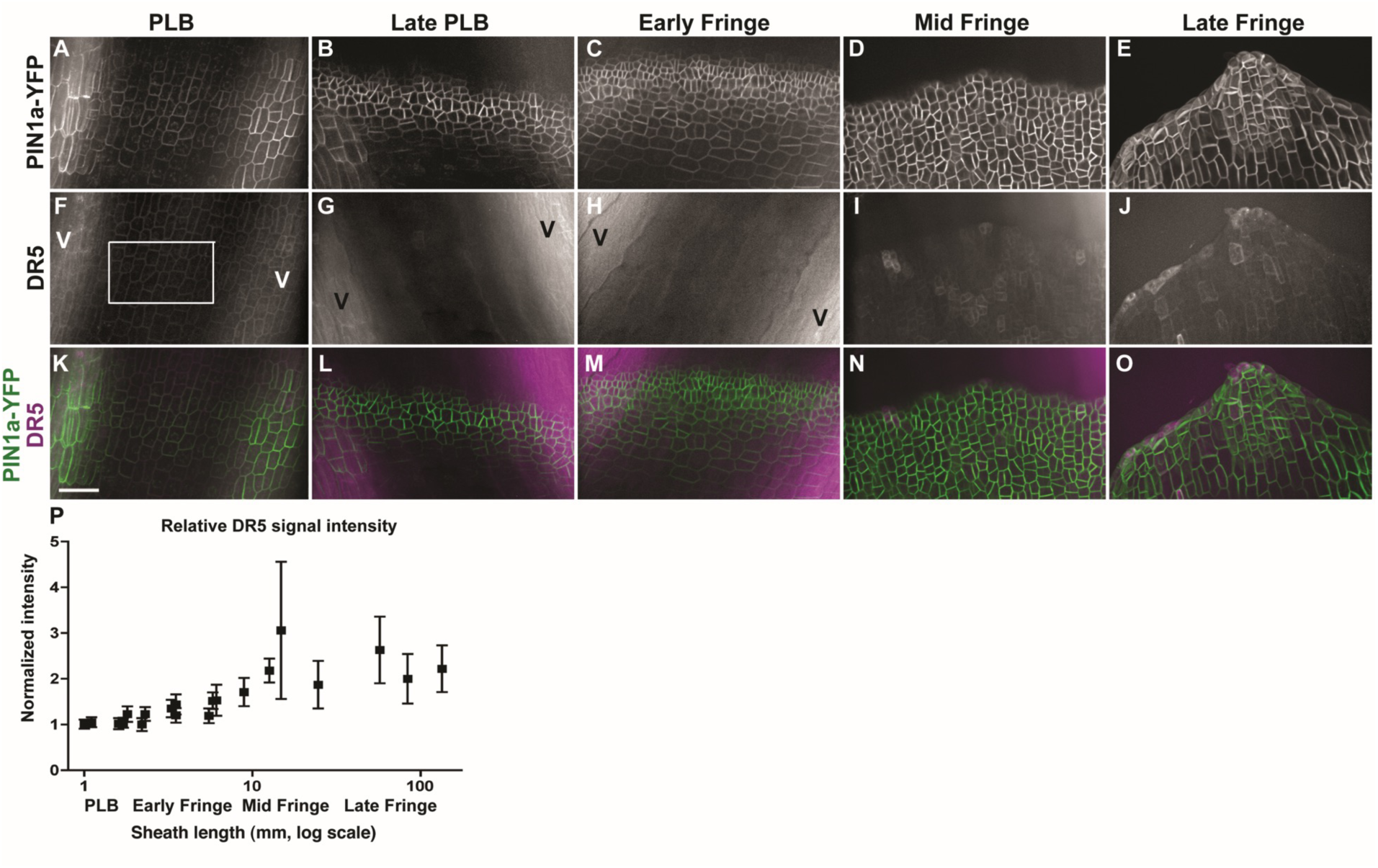
PIN1a-YFP and DR5rev:mRFPer (DR5) localization during ligule development. PIN1a-YFP expression is shown in the top panels (A-E); DR5 in the middle panel (F-J) with merged PIN1a-YFP (green) and DR5 (magenta) in the bottom panel (K-O). Normalized DR5 fluorescence intensity values are shown (P). Each point is the average of three or more measurements per sample. DR5 intensity values increased significantly and were more variable during later ligule stages (F-test, p<0.01). Error bars show standard error. Scale bar = 50 μm (all micrographs have the same magnification).

### PIN1a-YFP is less polarized in the PLB and ligule fringe compared to the blade

PIN proteins are auxin efflux transporters, and polar localization of PINs can result in directional auxin flow. PIN1 polarization, defined as asymmetric accumulation of the protein between polar domains of the plasma membrane, correlates with the direction of auxin transport (Wisniewska et al., 2006). We examined PIN1a-YFP localization in the developing ligule, as an indication of directional auxin transport. We compared the relative polarity of PIN1a-YFP in developing ligule cells to that of blade epidermal cells (Fig. 5K-M). In contrast to the blade, where clear PIN1a-YFP polarity was observed (Fig. 5J), PIN1a-YFP localization in PLB and ligule cells appeared relatively nonpolar. Consistent with this observation, the fluorescence intensity ratio of PIN1a-YFP in the blade was 1.71 +/- 0.11 showing that the PIN1a-YFP is polarized in the blade, as expected (Fig. 5L,M). PIN1a-YFP was primarily localized to the rootward side of blade epidermal cells consistent with basipetal auxin transport in leaf primordia (Sawchuk et al., 2013; Scarpella et al., 2010). In contrast, PIN1a-YFP signal was significantly less polarized in the PLB and the forming ligule (Fig. 5K,M), with mean ratios ranging from 1.21 +/- 0.04 to 1.47 +/-0.09 (Fig. 5M). Whereas PIN1a-YFP signal was strong in the PLB and ligule throughout development, its subcellular localization was relatively nonpolar.

### Auxin transcriptional responses reported by DR5 are low in the PLB and increase during ligule elongation

*DR5* is a synthetic promoter containing auxin response elements, which can be used in combination with reporters to approximate auxin transcriptional responses (Ulmasov et al., 1997). We examined expression of *DR5rev:mRFPer* (*DR5*) in plants coexpressing *PIN1a-YFP* (Fig. 6), to determine if changes in auxin responses correlated with proximal-distal specification in the PLB. If auxin transcriptional responses were associated with the specification of the ligule founder cells, high DR5 signal would be expected in the proximal and central regions of the PLB prior to ligule outgrowth, similar to the localization of PIN1a-YFP. DR5 signal was high in the underlying vasculature, which indicated auxin responses in those regions (Fig. 6). Special care was taken to measure DR5 intensity only in epidermal cells located between vascular bundles, thus excluding the strong signal from underlying vasculature (e.g. areas labeled with a V for vasculature in Fig. 6). In contrast to strong PIN1a-YFP signal (Fig. 6 A-E), DR5 signal was weak throughout the entire PLB, and gradually increased after the ligule fringe formed (F-test, p<0.01, Fig. 6F-P). These data suggest that DR5-related auxin responses do not underlie the specification of the ligule founder cells, although we cannot rule out the possibility that a distinct, DR5-independent subset of auxin responses may occur.

## Discussion

A key problem in plant development is understanding how new growth axes are generated distinct from pre-existing growth axes. Establishment of boundaries and boundary-like domains can help facilitate the physical separation of new organs or structures by locally limiting growth, but the mechanisms restricting cell expansion in boundaries are not clear (Bell et al., 2012; Gendron et al., 2012; Lee et al., 2009). In maize, the formation of the ligule is a particularly complex morphogenic process because a thin flap forms entirely from epidermal cells and cleanly diverges from the rest of the epidermis along a well-defined cleft at the ligule-auricle junction.

Here we examine cellular properties during early ligule development. In the early PLB (Fig. 7B), PIN1a-YFP localizes throughout the entire PLB, with the strongest signal overlying the vasculature. At this stage, cell division orientation is exclusively anticlinal, epidermal cell depth is uniform, and the topography of the leaf surface is nearly flat in the proximodistal direction. Epidermal cell walls are more rigid in the PLB compared to the blade and sheath. In the PLB stage (Fig. 7C), PIN1a-YFP signal becomes stronger and more uniform throughout the entire ligular region, but the protein remains relatively nonpolar at the subcellular level. During the PLB stage, cells in the sheath epidermis and proximal ligular region increase in depth considerably more than the distal ligular region and blade, and periclinal divisions are observed in the proximal ligular region. These changes contribute to the formation of a ridge that forms immediately proximal to the zone with the most rigid cell walls. During the late PLB stage (Fig. 7D), the zone of PIN1a-YFP accumulation narrows, localizing to the nascent ridge. The frequency of periclinal divisions reaches a maximum and the cells on the ridge have softer cell walls. At this stage, the proximal and distal zones of the ligular region differ in epidermal thickness, division plane orientation, cell wall rigidity, and PIN1a-YFP accumulation. In the early fringe stage, the soft cells on the more proximal ridge grow over the top of the more distal rigid cells, forming a well-defined cleft (Fig. 7E). The newly separated growth axis of the early ligule fringe then elongates primarily via transverse divisions and cell expansion. Our findings are consistent with the model proposed by Johnston et al,. (2014), which was based on expression profiling, that PLB is partitioned into subdomains prior to ligule outgrowth. Aside from preligule-preauricle specification, there may be additional subdomains that are not currently recognized.

**Figure 7:**
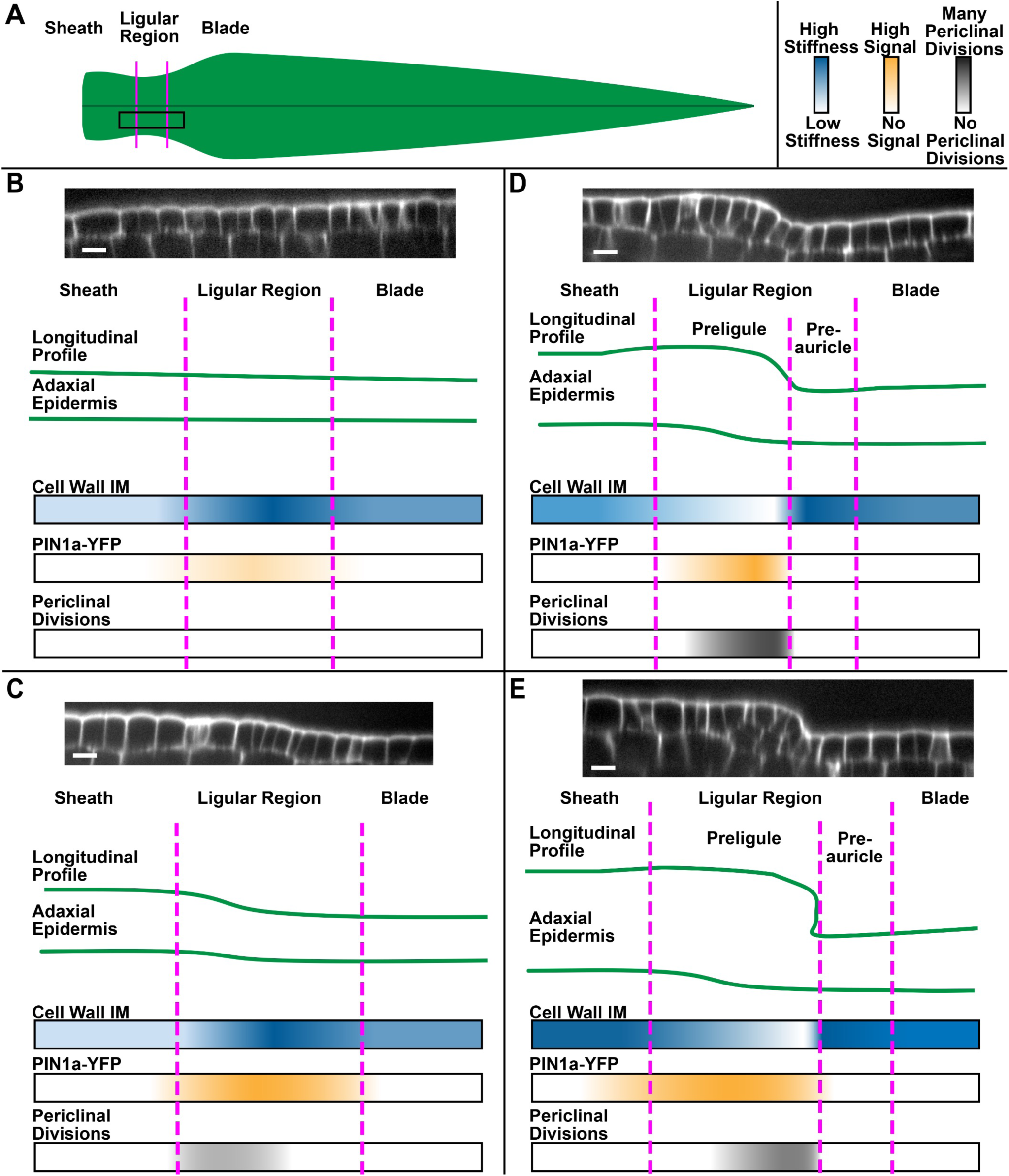
Summary of patterns observed during early ligule development. (A) Cartoon of maize leaf primordium and proximodistal zones. Black box indicates the area studied. (B-D) Patterns in epidermal topography are shown in longitudinal profile of the epidermal cell layer using representative confocal micrograph projections, and cartoons without individual cells. Proximodistal patterns in cell wall IM, PIN1a-YFP signal, and the position/frequency of periclinal divisions are shown using color gradients. (B) Early PLB stage, (C) PLB Stage, (D) Late PLB Stage, (E) Early Fringe Stage. Scale bars=15 μm.

The abrupt shift from anticlinal to periclinal division orientation is a key feature of developmental events in plants. Mechanisms regulating this shift are not well understood. Previous predictive modeling of cell divisions via soap-film minimization showed the geometry of cells in the late PLB favors periclinal divisions (Martinez et al., 2018). Late PLB cells are small in the epidermal surface, but relatively thick in the depth axis, resulting in a columnar cell shape. Cells tend to divide along the shortest axis, so the Martinez et al. (2018) geometry-based surface minimization model most commonly predicts periclinal divisions in these cells. Our data show that the earliest periclinal divisions in the PLB stage are observed in the proximal and central PLB cells, which have thickened more in the depth axis than the distal cells. Furthermore, the periclinal division rate is highest at the late PLB stage, when cell area at the epidermal surface is the smallest. Differential cell thickening establishes a geometry in the proximal and central PLB cells that favors periclinal division plane orientation.

Our data add to the existing body of nanoindentation and AFM experiments on live plant cells (Bou Daher et al., 2018; Majda et al., 2017; Peaucelle et al., 2011; Routier-Kierzkowska et al., 2012). The dramatic softening of cell walls in the proximal PLB preceding ligule outgrowth is highly reminiscent of data from the Arabidopsis SAM, where biochemical changes and mechanical softening in the cell walls of the subepidermal cell layers precede the outgrowth of leaf primordia (Peaucelle et al., 2011). The juxtaposition between rigid and soft epidermal cells along a discrete line is conspicuous, and suggests that differential regulation of cell wall properties within adjacent cell populations mechanically contributes to the sharp cleft at the preligule-preauricle junction, reminiscent of earlier studies showing that mechanical patterns contribute to abrupt changes in directional growth at the shoot apex (Selker et al., 1992). PLB cells also exhibit reduced elastic asymmetry between transverse and longitudinal wall segments. Elastic asymmetry was shown to correlate with anisotropic expansion in the Arabidopsis hypocotyl (Bou Daher et al., 2018). A shift to isotropic growth is observed during leaf initiation from the SAM peripheral zone (Sassi et al., 2014), so it is possible that a similar trend may occur during early ligule outgrowth. These findings may inform future experiments exploring differences in cell expansion and growth anisotropy during maize leaf development.

We note that the indentation modulus of the cell wall is not a direct indicator of extensibility nor actual cell expansion, which is a plastic, irreversible process (Cosgrove, 2016). The cell wall is heterogeneous and materially anisotropic, each cell exists within the structure of multiple tissue layers, and changes in wall chemistry, heterogeneity and degree of plasticity occur during growth and development. Computational modeling could help explore the mechanics of nanoindentation in live plant tissue, and the biological implications of the elastic properties of the cell wall. Finally, more experiments are necessary to determine the cell wall components, remodeling enzymes, or other properties underlying the observed differences in rigidity between epidermal regions.

The upregulation of PIN-like genes in the PLB has been previously reported, suggesting a role for auxin transport in ligule development (Johnston et al., 2014; Moon et al., 2013). Polar auxin transport and high auxin transcriptional responses are associated with the initiation and development of many structures during plant development, including leaves, branches, lateral roots, root hairs, and vasculature (Barazesh and McSteen, 2008; Bennett et al., 2014; Du and Scheres, 2018; Hajný et al., 2022; Jones et al., 2009; Leyser, 2018; McSteen and Leyser, 2005; Pitts et al., 1998; Scarpella et al., 2010). We observed that the PIN1a-YFP-expressing zone narrowed significantly during the late PLB stage, becoming restricted to the small PLB cells in the center of the ligular region. This could be consistent with the focusing of auxin toward a convergence point, as it is during leaf initiation (Conklin et al., 2019). However, we found that PIN1a-YFP accumulation in the PLB and ligule was not polarly localized, and DR5 signal was consistently low throughout the PLB prior to outgrowth of the ligule fringe. One possible interpretation is that auxin is being transported away from the PLB, locally depleting auxin and preventing *DR5* expression. This is puzzling because PIN1a is itself an auxin-responsive gene. In addition, other auxin-related genes, including several *AUXIN RESPONSE FACTOR*s (*ARF*s) and *SMALL AUXIN UP-REGULATED RNA*s (*SAUR*s), and *GRETCHEN HAGEN3* (*GH3*) genes are differentially expressed in the PLB (Johnston et al., 2014). *GH3* genes encode IAA-amino acid synthetases, which conjugate IAA to aspartate, rendering the auxin biologically inactive and marked for storage or degradation (Jez, 2022; Staswick et al., 2005). Nonpolar auxin efflux, increased catabolism or sequestration, and a lack of DR5 signal are consistent with low auxin transcriptional responses in the PLB, rather than the elevated responses associated with the initiation of many other plant organs. There are 15 *PIN* genes in maize (Yue et al., 2015), several of which are upregulated in the PLB, such as *PIN5, PIN1c,* and *SoPIN1/PIN1d* (GRMZM2G171702) (Johnston et al., 2014). Other PIN proteins could localize differently than our PIN1a-YFP construct. For example, AtPIN1 is involved in polar auxin transport in the epidermis of the Arabidopsis meristem during leaf initiation, but in maize this role is filled by SoPIN1/PIN1d, which belongs to the SISTER-OF-PIN1 (SoPIN1) clade, while AtPIN1 orthologs ZmPIN1a and ZmPIN1b act in internal tissue layers (Carraro et al., 2006; Li et al., 2019; O’Connor et al., 2014). The expression of *TIR1/AFB*s, *IAA*s, and *ARF*s, and differential affinities for auxin, can affect the sensitivity of auxin signaling in a given region (Vernoux et al., 2011). While *TIR/AFB* auxin receptor genes are expressed relatively consistently between the blade, ligular and sheath zones, both *ARF*s and *IAA*s are differentially expressed in the PLB (Johnston et al., 2014). It is possible that a distinct subset of auxin responses is activated in the PLB without high *DR5* expression. Particularly, *GRMZM2G158359*, a likely ortholog of the transmembrane noncanonical auxin receptor gene *AtTMK1*, is significantly upregulated in the PLB (FDR<0.05, Johnston et al., 2014), suggesting that extracellular auxin could serve a signaling role in the PLB without activating canonical TIR1/AFB-AuxIAA signaling (Cao et al., 2019; Lin et al., 2021; Xu et al., 2014). With so much complexity governing auxin signaling and responses, the role of auxin in the development of the ligular region remains unclear.

We demonstrate here that the boundary between blade and sheath in the maize leaf is progressively refined in the ligular region, producing two subdomains. Shifts in topography, cell growth, division orientation, and PIN localization correlate with changes in cell wall biophysical properties in the ligular region. The rigid PLB is partitioned into a soft, proximal incipient ligule, and a rigid, distal preauricle zone. These events correlate with ligule outgrowth and presage the development of the auricle between the ligule and blade. How this occurs across the three dimensions of the leaf is an intriguing question, given the auricle hinge becomes anatomically unique in all dimensions. Next steps are to refine molecular and cellular changes in the transverse and mediolateral three-dimensions to fully understand how ligule- and auricle-specific cell growth is coordinated.

## Methods

### Plant Growth and Dissection

Maize plants were grown in two-gallon pots in standard greenhouse conditions (28 °C, 16 hours light, 8 hours dark) for 2-4 weeks at the Laramie Research and Extension Center at the Agriculture Experiment Station at the University of Wyoming or in greenhouses under similar conditions at UC Riverside. Maize plants used for imaging included the inbreds B73, Mo17, and plants containing fluorescent markers developed by the Maize Cell Genomics project (http://maize.jcvi.org/cellgenomics/index.php). Maize lines expressing *PIN1a-YFP* and *DR5rev*:*mRFPer* (Gallavotti et al., 2008)*, DR5, YFP-TUBULIN, CFP-TUBULIN*, or *TAN-YFP* have been previously described (Mohanty et al., 2009). Transgenic plants were selected by resistance to a solution of 4g/L glufosinate-ammonium (Basta, Bayer Sciences) in 0.5% Tween, applied to the leaf. Plants were genotyped by PCR using primers CYFP LSP1 (5’ agcgcgatcacatggtcct) and PIN4110R (5’ ttcccgaagctgaagtcgtcc) or DR5-870F (5’ tgaagggcgagatcaagatgag) and DR5-1225R (5’ ctcaacacatgagcgaaacc).

For dissections, leaves were sequentially removed from the plant and leaf numbers counted from leaf 1 in toward the SAM. The length of the sheath region was measured with calipers, and the ligule growth stage was assessed by either confocal or scanning electron microscopy (all stages), or in a separate set of experiments by atomic force microscopy (AFM) (below 3.5mm sheath length). The leaves examined ranged from leaf number 4 to 12, depending on plant age and the developmental stage at which the plants were collected.

### Imaging and measuring cell size and division arrays using YFP-TUBULIN lines

Adaxial ligule regions of freshly dissected plants were mounted in water in Rose chambers and micrographs were analyzed for cell area and division plane orientation using ImageJ (http://rsbweb.nih.gov/ij/). Angles of preprophase bands or phragmoplasts were classified as anticlinal transverse, anticlinal longitudinal, periclinal, or oblique relative to the long axis of the leaf, as shown in Fig. 2C-F.

At the PLB and late PLB stages, the relative position of actively dividing cells within the PLB was determined using ImageJ. First, the proximal and distal extremities of the ligular region were traced according to differences in cell size and shape. Then, actively dividing cells, as indicated by the presence of a preprophase band labeled with either YFP-TUBULIN or TAN-YFP, were located within the PLB. Their relative position was calculated by measuring the distance from the proximal end of the PLB to the center of the dividing cell, and then from the proximal end to the distal end of the PLB, and dividing the former value by the latter (Fig. S3). This generates values ranging from 0 at the proximal end of the PLB to 1 at the distal end.

### Confocal microscopy

Images were acquired on two spinning disk confocal microscopes. The EM-CCD camera (ImagEM, Hamamatsu) was mounted on an IX71 stand equipped with a spinning-disc confocal head (CSU-X1; Yokogawa). A LMM5 laser launch was used to provide illumination (Spectral Applied research). Laser lines of 488 and 561 nm were used to excite PIN1a-YFP, YFP-TUBULIN, TAN1-YFP and *DR5rev*:*mRFPer* with band pass filters ET525/50M and ET595/50M (Chroma Technology) respectively. Some image acquisition was performed using Metamorph 7.7 software (Molecular Devices). Images were acquired using 20x (0.85 NA) and 40x oil (1.30 NA) Olympus objectives. For additional samples in the early PLB, PLB, and late PLB stages of ligule development, the dissected ligular region was stained with 10 μg/mL propidium iodide for 10 minutes and mounted in water. Confocal scans were collected using the 40X objective lens through the epidermis with a Z-step of 0.2 μm using a Hamamatsu 9100C EM-CCD camera mounted on a Nikon Ti stand with a spinning disc confocal head (CSU-W1, Yokogawa) and a 40X water (1.1 NA) Nikon water objective. Propidium iodide, TAN1-YFP or YFP-TUBULIN, and/or CFP-TUBULIN were excited at 561nm; 514nm; 445nm and collected at 620/20nm; 540/30nm; 480/40 nm respectively.

### Scanning electron microscopy (SEM)

Two-, three- and four-week-old B73 leaf samples were used for SEM to characterize ligule stages. Sheath lengths were measured and the ligular region was excised with a scalpel, mounted with two-sided tape and loaded directly into the sample chamber of the tabletop electron microscope (Hitachi TM-1000), with included software used to acquire images.

### Image and statistical analysis

Image analysis was performed using ImageJ (*rsbweb.nih.gov/ij/*), FIJI (ImageJ), or Metamorph v. 7.7. Data were analyzed in Excel and Access (Microsoft Office) and graphs produced in Graphpad (Prism) and R. For measuring PIN1a-YFP polarity (Fig. 5M), 30 cells at each stage from four different plants were used for analysis. PIN1a-YFP fluorescence intensity measurements were performed by scanning through a Z stack of an entire epidermal cell. The plane with the highest fluorescence intensity value was selected at each side of the randomly selected cell. A one-pixel thick line was drawn across each side of the cell cortex and average intensity values were recorded. Ratios were calculated by dividing the highest average intensity value by the lowest for each analyzed cell in Excel and the error bars are 95% confidence intervals.

For measuring DR5 fluorescence intensity, a box was drawn between veins to avoid fluorescence of underlying vasculature (see Fig. 6B). Average fluorescence intensity was recorded from the boxed area of at least 1000 square pixels from a five µm deep maximum projection. For DR5 expression, three plants were used per stage and standard error bars are shown in Fig. 4. The average intensity values from areas between vascular bundles of three samples in a single plant were normalized by dividing each value by the lowest average DR5 intensity value. Significance tests comparing the distribution of PIN1a-YFP fluorescence intensity ratios were performed using the non-parametric Kolmogorov-Smirnov (KS) test.

Cell depth was measured for PI-stained leaves dissected from three plants in the early PLB, PLB, and late PLB stages using ImageJ. The image stack was projected as an orthoslice and the thicknesses of cells were measured at transverse wall segments by counting the number of z-steps between the top and bottom of the wall segment. Statistical differences were assessed via the non-parametric Kruskal-Wallis test, with a Dunn’s post-hoc for pairwise comparisons.

When analyzing AFM data, the difference in sheath length between the two observed tissue-level mechanical patterns was assessed via a Mann-Whitney U-test. For manual resampling, at least 50 indentations were used per wall category per tissue zone per sample to calculate average IM values. After manual resampling, global variation in IM with respect to wall category and tissue zone were assessed via Kruskall-Wallis tests. Then pairwise Wilcoxon signed rank tests were performed at significance levels of p<0.05 and p<0.01, using the *W*-statistic. Variation in IM with respect to developmental stage was assessed via Mann-Whitney U-tests.

### Atomic Force Microscopy

Developing ligules were dissected as described above and sheath length was measured using electronic calipers. The samples were quickly placed in 0.55M mannitol solution for at least 15 minutes to induce plasmolysis, before being affixed to a microscope slide using double-sided tape. Additional mannitol solution was used to immerse the sample and pre-wet the probe.

AFM was performed using a JPK Nanowizard 4a AFM in force mapping mode, at the California NanoSystems Institute at UCLA. Indentations were performed with a constant maximum force of 500 nN, with extend and retract times of 0.1s, at a spatial resolution of at least one indentation per 2 microns. The probes used were PPP-NCL probes with a 10 nm pyramidal tip, with an average force modulus of 45 N/m Each tip was calibrated separately to accommodate slight differences.

Data were processed using the JPKSPM software. The raw indentation data were converted into IM (indentation modulus) using the Hertzian contact model as described (Peaucelle et al., 2011). Because the maximum scan area was too small to adequately sample all epidermal regions in a single scan, multiple overlapping scans were measured, processed, and manually reassembled by identifying cell walls in the overlapping areas. To quantify IM along the longitudinal axis of the leaf, regional IM values were averaged using a sliding window approach. For a given leaf, a 25 μm-wide rectangle was drawn, and repositioned along the longitudinal axis until the local maximum for average IM was located in the PLB. This position was designated relative position 0, and the average IM for that bin was set as 1. Regional averages for IM were then measured along the proximodistal axis in 25 μm-wide bins, with a 12.5 μm step between bins. Position and average IM were normalized to the local maximum at the PLB for each leaf measured. Manual resampling of the scans was performed using a custom script in MATLAB (github.com/mathworks). Force maps were projected as a heatmap and at least 50 pixels within each epidermal zone and cell wall category were selected and averaged for each sample.

## Acknowledgements

We thank Dr. Holly Steinkraus at the University of Wyoming (UW) for assistance and support of the research, the UW Molecular and Life Sciences Graduate Program, Dr. Jay Gatlin for use of the spinning disc microscope and resources in his laboratory at UW, the California NanoSystems Institute for AFM access, and Dr. Pablo Martinez for advice and training on maize dissections. We acknowledge grant funding from the NSF IOS-1052051 and DBI 0501862 to AWS, NSF-Career 1942734, NSF 1716972, and NSF1244202 to CGR, USDA-AFRI 2012-67014-19429 and USDA CA-R-BPS-7659-H to PSS, NSF1548571 Science and Technology Center for Engineering Mechano-Biology to SB, and US Department of Education GAANN grant P200A210029, for partial support of WN.

## Supplementary figure legends

**Figure S1:**
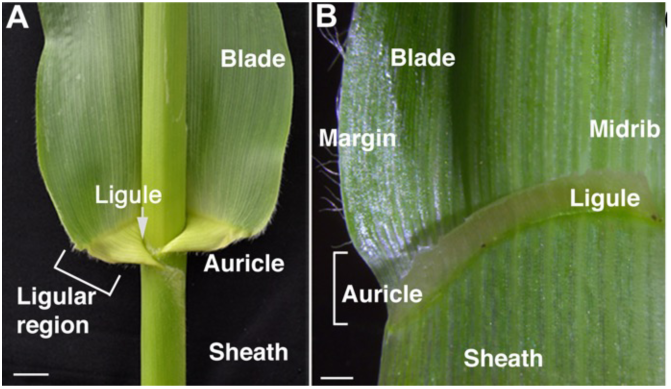
Maize leaf structure. The maize leaf is composed of a distal blade and proximal sheath separated at the ligular region (bracket), consisting of a ligule and auricle. Scale bar = 2 cm. (B) The leaf is cut at the midrib to expose the adaxial view of the ligular region. Scale bar = 1 mm.

**Figure S2:**
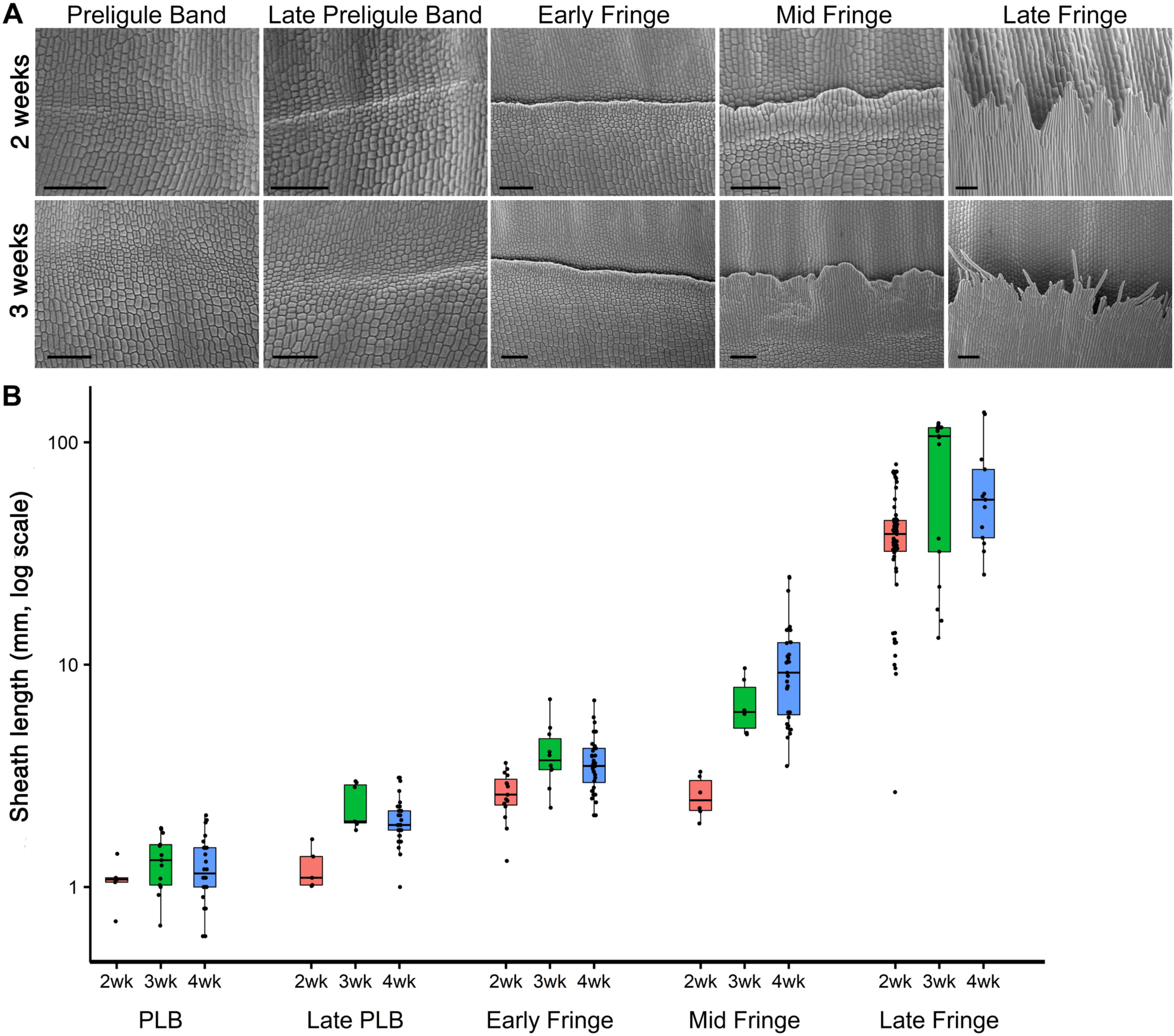
Stages of ligule development relative to sheath height in 2-, 3-, and 4-week-old plants. Scanning electron micrographs of sequentially dissected leaves of 2- and 3-week-old maize plants. (B) Stages of ligule development in 2-, 3-, and 4-week-old maize plants relative to sheath length. Scale bar = 100 μm.

**Figure S3:**
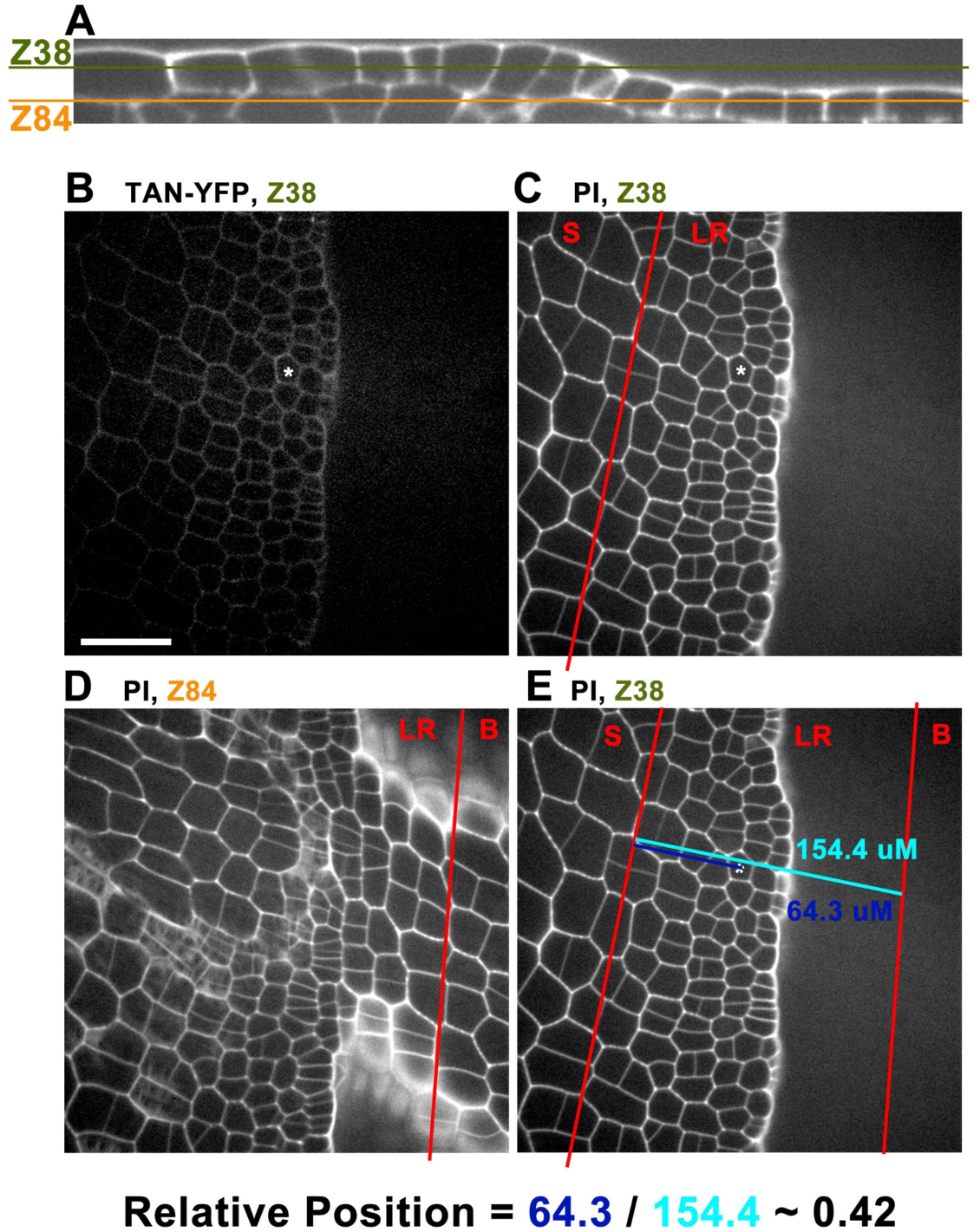
Cell division relative position calculation example. Example of how relative position within the ligular region was calculated for dividing cells. (A) Orthogonal projection of confocal scan through preligule epidermis showing position of Z38 and Z84. B,C, and E are at Z38, in plane with the sheath, and D is at Z84, in plane with the blade. (B) Periclinal divisions were identified using either TAN-YFP and propidium iodide (shown here), or YFP-Tubulin. Asterisk indicates periclinally dividing cell. (C,D) Estimated position of proximal and distal extremities of the ligule region identified from PI-stained cell outlines. Ligular region identified by differences in cell size and shape from adjacent sheath and blade cells. (E) Teal line indicates length of ligular region at that mediolateral position. Fuschia indicates distance from proximal extremity of ligular region to the center of the marked cell. Scale bar = 50 μm, all images same scale.

**Figure S4:**
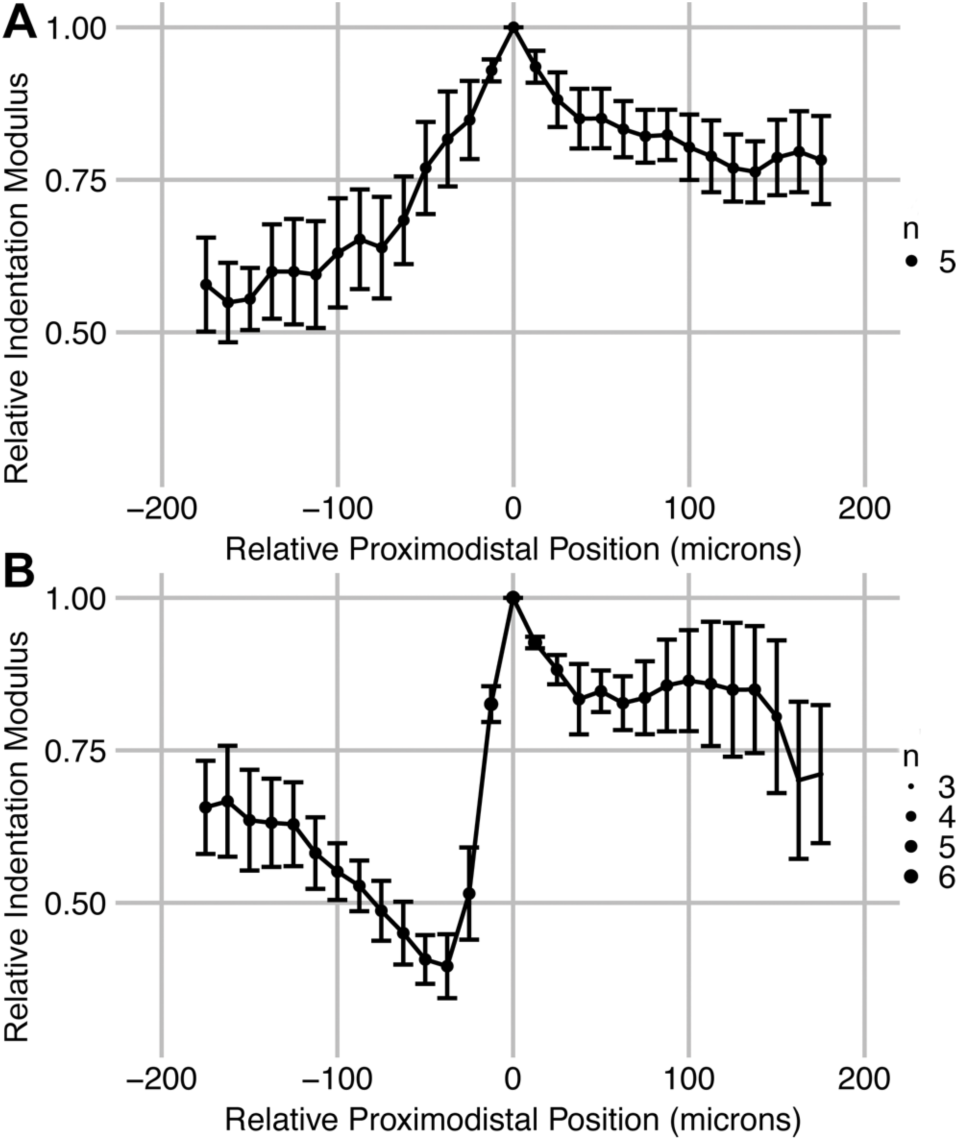
AFM force mapping sliding window analysis of Mo17 leaves. Sliding window regional averages in IM. Data normalized to the local maximum in the PLB, set as position=0, relative IM=1. Top panel is the average of all Mo17 plants (n=5) exhibiting the early mechanical pattern. Bottom panel is the average of all Mo17 plants (n=6) exhibiting the late mechanical pattern. Error bars indicate standard error.

**Figure S5:**
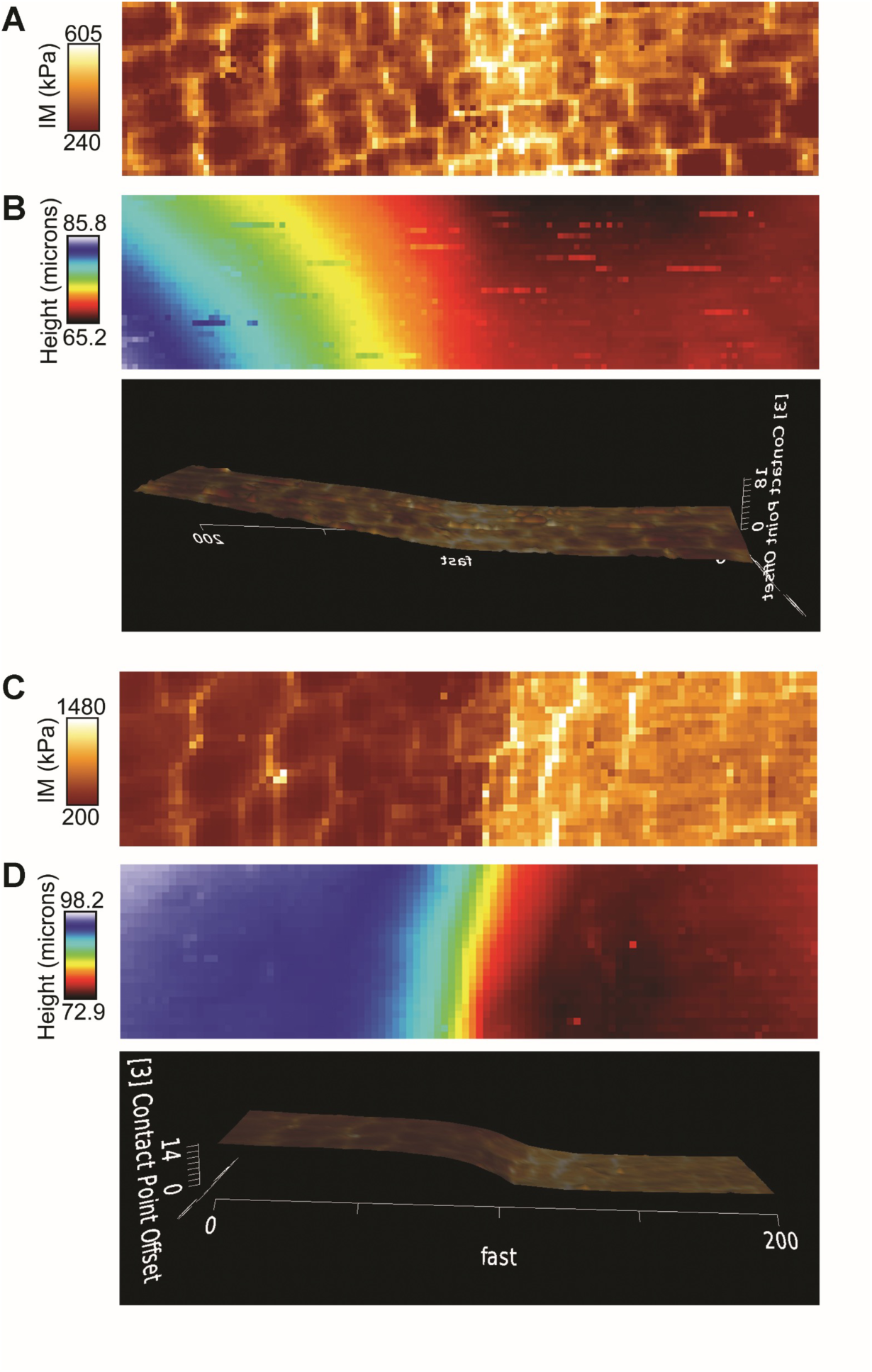
Mechanical transition within PLB relative to topography. (A) AFM scan of a leaf in the PLB stage, sheath length 0.6mm. Resolution is 1.6 μm per pixel. (B) Topography of the same sample as (A). This height map was first normalized along the mediolateral axis to highlight topography in the proximodistal direction. Height portrayed via both heatmap and 3D projection (C) AFM scan of a leaf in the late PLB stage, sheath length 2.0mm. Resolution is 2.0 μm per pixel. (D) Topography of the same sample as (C). Scale bars = 50 μm.

**Figure S6:**
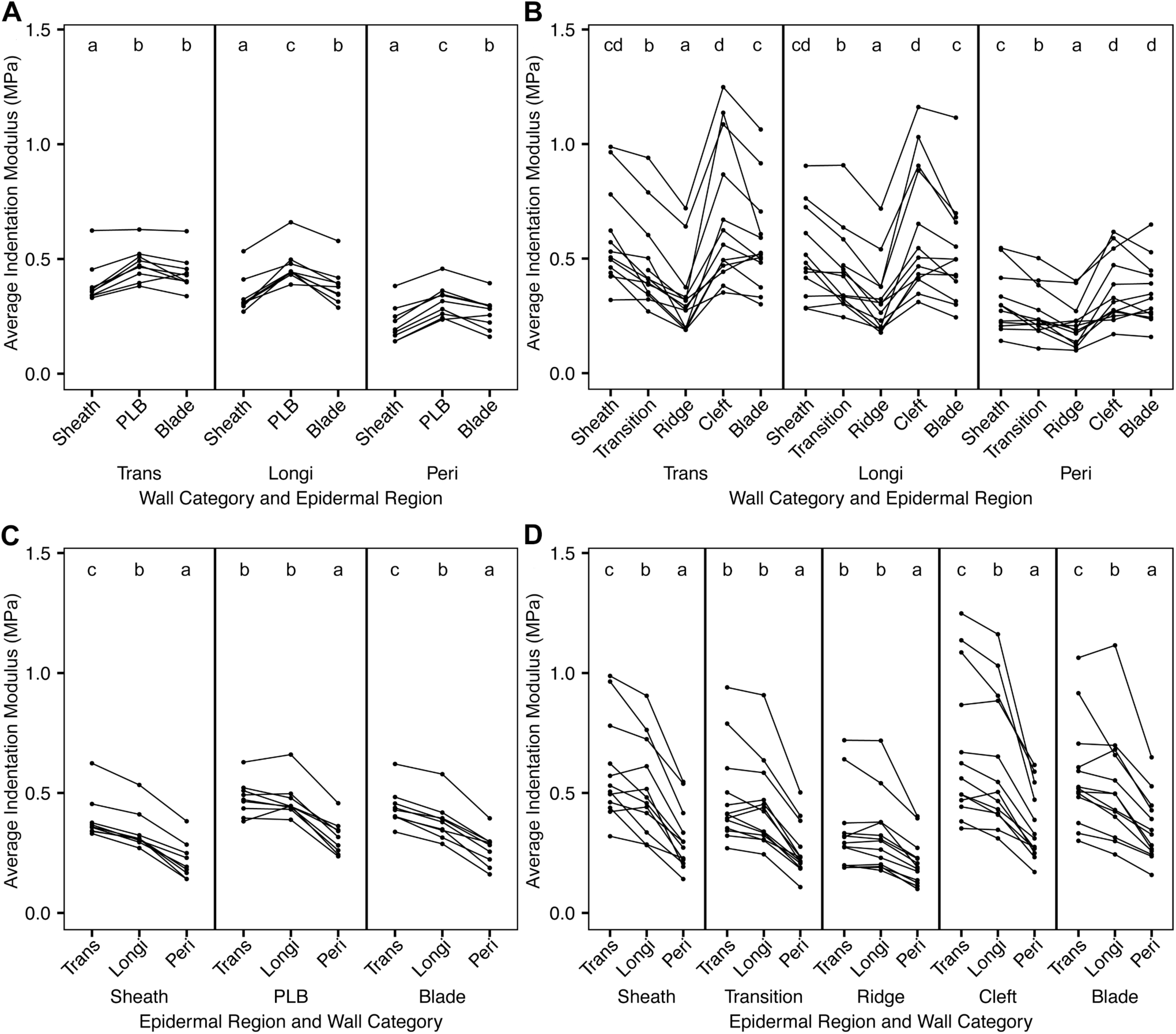
Manual resampling reveals regional and subcellular patterns in IM. IM was resampled for each wall category in each epidermal region of each sample. Samplings from different wall categories or tissue zones in the same leaf are connected with lines. Each dot indicates the average IM of at least 50 indentations from a particular wall category in a particular epidermal zone in one leaf. (A) All early-stage B73 samples, grouped by epidermal zone. (B) All late-stage B73 samples, grouped by epidermal zone. (C,D) Same data but instead grouped by wall category. Statistical analysis was performed independently for each panel and subpanel. Statistical significance was determined via Kruskall-Wallis test followed by pairwise Wilcoxon signed rank tests using the *W*-value at an alpha of p<0.05.

---

**Table S1:** Cell division orientation during ligule developmental stages. 3-5 leaves expressing YFP-Tubulin were imaged per stage. To calculate cell area in each leaf, three boxes each encompassing 20-100 cells were drawn spanning the PLB or over a portion of the elongating ligule in the confocal micrographs, and the number of cells in each box was counted. Areas of the boxes were divided by the number of cells in each box to calculate the average cell area. To calculate cell division orientation %, the number of mitotic cells in each orientation was counted in each sample, and divided by the total number of mitotic cells (11-46 mitotic cells imaged per sample).

| Developmental stage | n<br>Individual leaves measured per stage | Sheath length range in mm | Cell size ($\mu\text{m}^2$ +/- S.E.) | Total mitotic cells observed | Anticlinal longitudinal division (% +/- S.E.) | Anticlinal transverse divisions (% +/- S.E. ) | Periclinal divisions (% +/- S.E. ) | Oblique divisions (% +/- S.E.) |
| --- | --- | --- | --- | --- | --- | --- | --- | --- |
| Early Preligule band | 4 | 0.8 - 1.3 | 158 +/- 9<br>n=414 | 77 | **52 +/- 9%** | 49 +/- 8% | 0% | 3 +/- 2% |
| Preligule band | 3 | 1.5 - 2.1 | 134 +/- 9<br>n=315 | 59 | 38 +/- 3% | **45 +/- 5%** | 10 +/- 4% | 5 +/- 3% |
| Late Preligule band | 5 | 1.8 - 3.1 | 105 +/- 6<br>n=759 | 132 | 18 +/- 2% | 31 +/- 2% | **46 +/- 1%** | 4 +/- 2% |
| Early fringe | 3 | 2.5 - 4.4 | 186 +/- 16<br>n=304 | 62 | 22 +/- 1% | **48 +/- 1%** | 27 +/- 2% | 2 +/- 2% |
| Mid fringe | 3 | 5.4 - 12.5 | 450 +/- 145<br>n=539 | 45 | 9 +/- 5% | **71 +/- 9%** | 16 +/- 5% | 2 +/- 2% |
| Late fringe | 3 | 35.1 - 57.5 | 1511 +/- 147<br>n=246 | 0 | n/a | n/a | n/a | n/a |

**Table S2:** Pairwise comparisons of average IM between epidermal zones via Wilcoxon signed rank test using the W-statistic.

| Comparison<br>(Epidermal Tissue Zones) | Wall Category | Average IM Zone 1 +/- s.d. (MPa) | Average IM Zone 2 +/- s.d. (MPa) | *W*-value | Significance level |
| --- | --- | --- | --- | --- | --- |
| Early Sheath v. Early PLB | Trans | 0.394 +/- 0.093 | 0.478 +/- 0.074 | 0 | p<0.01 |
|  | Longi | 0.340 +/- 0.082 | 0.470 +/- 0.077 | 0 | p<0.01 |
|  | Peri | 0.219 +/- 0.078 | 0.316 +/- 0.070 | 0 | p<0.01 |
| Early Sheath v. Early Blade | Trans | 0.394 +/- 0.093 | 0.441 +/- 0.079 | 1 | p<0.01 |
|  | Longi | 0.340 +/- 0.082 | 0.384 +/- 0.084 | 0 | p<0.01 |
|  | Peri | 0.219 +/- 0.078 | 0.265 +/- 0.069 | 0 | p<0.01 |
| Early PLB v. Early Blade | Trans | 0.478 +/- 0.074 | 0.441 +/- 0.079 | 6 | NS |
|  | Longi | 0.470 +/- 0.077 | 0.384 +/- 0.084 | 0 | p<0.01 |
|  | Peri | 0.316 +/- 0.070 | 0.265 +/- 0.069 | 1 | p<0.01 |
| Late Sheath v. Late Transition | Trans | 0.591 +/- 0.213 | 0.475 +/- 0.194 | 1 | p<0.01 |
|  | Longi | 0.518 +/- 0.196 | 0.443 +/- 0.180 | 9 | p<0.05 |
|  | Peri | 0.308 +/- 0.131 | 0.260 +/- 0.107 | 5 | p<0.01 |
| Late Sheath v. Late Ridge | Trans | 0.591 +/- 0.213 | 0.332 +/- 0.167 | 0 | p<0.01 |
|  | Longi | 0.518 +/- 0.196 | 0.323 +/- 0.156 | 0 | p<0.01 |
|  | Peri | 0.308 +/- 0.131 | 0.211 +/- 0.097 | 7 | p<0.01 |
| Late Sheath v. Late Cleft | Trans | 0.591 +/- 0.213 | 0.679 +/- 0.304 | 22 | NS |
|  | Longi | 0.518 +/- 0.196 | 0.620 +/- 0.281 | 18 | NS |
|  | Peri | 0.308 +/- 0.131 | 0.362 +/- 0.146 | 13 | p<0.05 |
| Late Sheath v. Late Blade | Trans | 0.591 +/- 0.213 | 0.571 +/- 0.218 | 25 | NS |
|  | Longi | 0.518 +/- 0.196 | 0.524 +/- 0.230 | 34 | NS |
|  | Peri | 0.308 +/- 0.131 | 0.351 +/- 0.134 | 13 | p<0.05 |
| Late Transition v. Late Ridge | Trans | 0.475 +/- 0.194 | 0.332 +/- 0.167 | 0 | p<0.01 |
|  | Longi | 0.443 +/- 0.180 | 0.323 +/- 0.156 | 0 | p<0.01 |
|  | Peri | 0.260 +/- 0.107 | 0.211 +/- 0.097 | 9 | p<0.01 |
| Late Transition v.<br>Late Cleft | Trans | 0.475 +/- 0.194 | 0.679 +/- 0.304 | 0 | p<0.01 |
|  | Longi | 0.443 +/- 0.180 | 0.620 +/- 0.281 | 0 | p<0.01 |
|  | Peri | 0.260 +/- 0.107 | 0.362 +/- 0.146 | 1 | p<0.01 |
| Late Transition v.<br>Late Blade | Trans | 0.475 +/- 0.194 | 0.571 +/- 0.218 | 0 | p<0.01 |
|  | Longi | 0.443 +/- 0.180 | 0.524 +/- 0.230 | 0 | p<0.01 |
|  | Peri | 0.260 +/- 0.107 | 0.351 +/- 0.134 | 0 | p<0.01 |
| Late Ridge v. Late<br>Cleft | Trans | 0.332 +/- 0.167 | 0.679 +/- 0.304 | 0 | p<0.01 |
|  | Longi | 0.323 +/- 0.156 | 0.620 +/- 0.281 | 11 | p<0.05 |
|  | Peri | 0.211 +/- 0.097 | 0.362 +/- 0.146 | 0 | p<0.01 |
| Late Ridge v. Late<br>Blade | Trans | 0.332 +/- 0.167 | 0.571 +/- 0.218 | 0 | p<0.01 |
|  | Longi | 0.323 +/- 0.156 | 0.524 +/- 0.230 | 0 | p<0.01 |
|  | Peri | 0.211 +/- 0.097 | 0.351 +/- 0.134 | 0 | p<0.01 |
| Late Cleft v. Late<br>Blade | Trans | 0.679 +/- 0.304 | 0.571 +/- 0.218 | 12 | p<0.05 |
|  | Longi | 0.620 +/- 0.281 | 0.524 +/- 0.230 | 7 | p<0.01 |
|  | Peri | 0.362 +/- 0.146 | 0.351 +/- 0.134 | 38 | NS |

**Table S3:** Pairwise comparisons of average IM between wall categories via Wilcoxon signed rank test using the *W*-statistic.

| Comparison<br>(Wall Categories) | Tissue<br>Zone | Category 1 Average IM<br>+/- s.d. (MPa) | Category 2 Average IM<br>+/- s.d. (MPa) | *W*-<br>value | Significance<br>level |
| --- | --- | --- | --- | --- | --- |
| Early Timepoint,<br>Trans v. Longi | Sheath | 0.394 +/- 0.093 | 0.340 +/- 0.082 | 0 | p<0.01 |
|  | PLB | 0.478 +/- 0.074 | 0.470 +/- 0.077 | 15 | NS |
|  | Blade | 0.441 +/- 0.079 | 0.384 +/- 0.084 | 0 | p<0.01 |
| Early Timepoint,<br>Trans v. Peri | Sheath | 0.394 +/- 0.093 | 0.219 +/- 0.078 | 0 | p<0.01 |
|  | PLB | 0.478 +/- 0.074 | 0.316 +/- 0.070 | 0 | p<0.01 |
|  | Blade | 0.441 +/- 0.079 | 0.265 +/- 0.069 | 0 | p<0.01 |
| Early Timepoint,<br>Longi v. Peri | Sheath | 0.340 +/- 0.082 | 0.219 +/- 0.078 | 0 | p<0.01 |
|  | PLB | 0.470 +/- 0.077 | 0.316 +/- 0.070 | 0 | p<0.01 |
|  | Blade | 0.384 +/- 0.084 | 0.265 +/- 0.069 | 0 | p<0.01 |
| Late Timepoint,<br>Trans v. Longi | Sheath | 0.591 +/- 0.213 | 0.518 +/- 0.196 | 0 | p<0.01 |
|  | Transition | 0.475 +/- 0.194 | 0.443 +/- 0.180 | 19 | NS |
|  | Ridge | 0.332 +/- 0.167 | 0.323 +/- 0.156 | 31 | NS |
|  | Cleft | 0.679 +/- 0.304 | 0.620 +/- 0.281 | 7 | p<0.01 |
|  | Blade | 0.571 +/- 0.218 | 0.524 +/- 0.230 | 16 | p<0.05 |
| Late Timepoint,<br>Trans v. Peri | Sheath | 0.591 +/- 0.213 | 0.308 +/- 0.131 | 0 | p<0.01 |
|  | Transition | 0.475 +/- 0.194 | 0.260 +/- 0.107 | 0 | p<0.01 |
|  | Ridge | 0.332 +/- 0.167 | 0.211 +/- 0.097 | 0 | p<0.01 |
|  | Cleft | 0.679 +/- 0.304 | 0.362 +/- 0.146 | 0 | p<0.01 |
|  | Blade | 0.571 +/- 0.218 | 0.351 +/- 0.134 | 0 | p<0.01 |
| Late Timepoint, Longi<br>v. Peri | Sheath | 0.518 +/- 0.196 | 0.308 +/- 0.131 | 0 | p<0.01 |
|  | Transition | 0.443 +/- 0.180 | 0.260 +/- 0.107 | 0 | p<0.01 |
|  | Ridge | 0.323 +/- 0.156 | 0.211 +/- 0.097 | 0 | p<0.01 |
|  | Cleft | 0.620 +/- 0.281 | 0.362 +/- 0.146 | 0 | p<0.01 |
|  | Blade | 0.524 +/- 0.230 | 0.351 +/- 0.134 | 0 | p<0.01 |

**Table S4:**
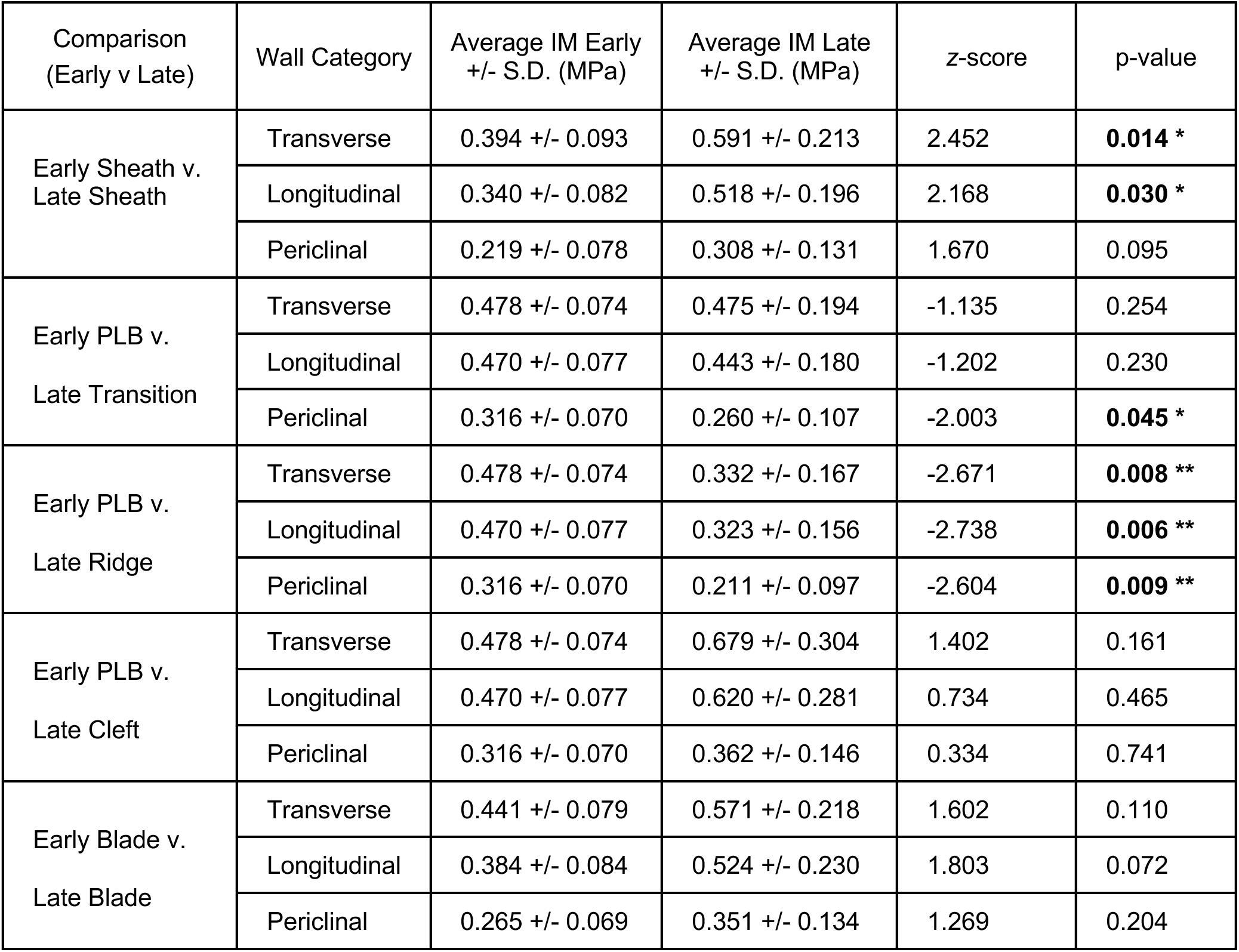
Pairwise comparisons of average IM between early and late mechanical stages via Mann-Whitney U-Test.

| Comparison<br>(Early v Late) | Wall Category | Average IM Early<br>+/- S.D. (MPa) | Average IM Late<br>+/- S.D. (MPa) | z-score | p-value |
| --- | --- | --- | --- | --- | --- |
| Early Sheath v.<br>Late Sheath | Transverse | 0.394 +/- 0.093 | 0.591 +/- 0.213 | 2.452 | **0.014 \*** |
|  | Longitudinal | 0.340 +/- 0.082 | 0.518 +/- 0.196 | 2.168 | **0.030 \*** |
|  | Periclinal | 0.219 +/- 0.078 | 0.308 +/- 0.131 | 1.670 | 0.095 |
| Early PLB v.<br>Late Transition | Transverse | 0.478 +/- 0.074 | 0.475 +/- 0.194 | -1.135 | 0.254 |
|  | Longitudinal | 0.470 +/- 0.077 | 0.443 +/- 0.180 | -1.202 | 0.230 |
|  | Periclinal | 0.316 +/- 0.070 | 0.260 +/- 0.107 | -2.003 | **0.045 \*** |
| Early PLB v.<br>Late Ridge | Transverse | 0.478 +/- 0.074 | 0.332 +/- 0.167 | -2.671 | **0.008 \*\*** |
|  | Longitudinal | 0.470 +/- 0.077 | 0.323 +/- 0.156 | -2.738 | **0.006 \*\*** |
|  | Periclinal | 0.316 +/- 0.070 | 0.211 +/- 0.097 | -2.604 | **0.009 \*\*** |
| Early PLB v.<br>Late Cleft | Transverse | 0.478 +/- 0.074 | 0.679 +/- 0.304 | 1.402 | 0.161 |
|  | Longitudinal | 0.470 +/- 0.077 | 0.620 +/- 0.281 | 0.734 | 0.465 |
|  | Periclinal | 0.316 +/- 0.070 | 0.362 +/- 0.146 | 0.334 | 0.741 |
| Early Blade v.<br>Late Blade | Transverse | 0.441 +/- 0.079 | 0.571 +/- 0.218 | 1.602 | 0.110 |
|  | Longitudinal | 0.384 +/- 0.084 | 0.524 +/- 0.230 | 1.803 | 0.072 |
|  | Periclinal | 0.265 +/- 0.069 | 0.351 +/- 0.134 | 1.269 | 0.204 |

